# Click-free imaging of carbohydrate trafficking in live cells using an azido photothermal probe

**DOI:** 10.1101/2024.03.08.584185

**Authors:** Qing Xia, Harini A. Perera, Rylie Bolarinho, Zeke A. Piskulich, Zhongyue Guo, Jiaze Yin, Hongjian He, Mingsheng Li, Xiaowei Ge, Qiang Cui, Olof Ramström, Mingdi Yan, Ji-Xin Cheng

**Author notes:** Correspondence (J.X.C.), (M.Y.).

## Abstract

Real-time tracking of intracellular carbohydrates remains challenging. While click chemistry allows bio-orthogonal tagging with fluorescent probes, the reaction permanently alters the target molecule and only allows a single snapshot. Here, we demonstrate click-free mid-infrared photothermal (MIP) imaging of azide-tagged carbohydrates in live cells. Leveraging the micromolar detection sensitivity for 6-azido-trehalose (TreAz) and the 300-nm spatial resolution of MIP imaging, the trehalose recycling pathway in single mycobacteria, from cytoplasmic uptake to membrane localization, is directly visualized. A peak shift of azide in MIP spectrum further uncovers interactions between TreAz and intracellular protein. MIP mapping of unreacted azide after click reaction reveals click chemistry heterogeneity within a bacterium. Broader applications of azido photothermal probes to visualize the initial steps of the Leloir pathway in yeasts and the newly synthesized glycans in mammalian cells are demonstrated.

## Introduction

Carbohydrates play important roles in various physiological processes. Most cell surfaces are coated with a layer of glycoproteins and glycolipids derived from post-translational modification inside the cell^1, 2^. Hence, monitoring intracellular trafficking of carbohydrates could elucidate the biological functions of carbohydrates in complex biological systems. Chromatography and mass spectrometry are used to quantify the amount of intracellular carbohydrates^3, 4^. Yet, these in vitro methods lack spatial information and their destructive nature prohibits live cell analysis. Fluorescence microscopy has been an invaluable tool for mapping intracellular carbohydrates with high selectivity. Accelerated efforts have been devoted to developing fluorescent analogs, based on fluorophores like boron-dipyrromethene and fluorescein, to track specific carbohydrates^5, 6^, including glucose^7^, maltose^8^ and trehalose^9^. However, these fluorophores typically have a size comparable to or larger than the target molecules^10^, which often alters the intracellular localization and uptake pathways of the target molecules. For example, a widely used fluorescent analog of glucose, NBD-glucose, has been shown to enter cells independent of the glucose transporter^7^.

To track molecules in their natural state, click chemistry has been developed for visualizing carbohydrates in live cells with minimal disruption to the surrounding cellular environment^11, 12, 13^. This technique relies on tagging of carbohydrates with an azido group and subsequent mapping of glycans with the conjugation of azide to an alkyne-tagged fluorophore via click reaction^14, 15^. The azide reporter, being small in size, has been shown not to interfere with cellular metabolism and has been used to study the function of carbohydrates in living systems, such as cell signaling^16^, immunomodulation^17^, and carbohydrate core formation^18^. However, the fluorophores commonly used in click chemistry exhibit a relatively low cell permeability^19^, thus requiring long incubation with biological samples ranging from 30 to 120 minutes and hindering real-time tracking of carbohydrates within living cells^14^. Photobleaching and cytotoxicity present additional challenges. Moreover, the click reaction efficiency inside the cell remains uncertain, potentially leading to inaccurate imaging results.

Here, we present a click-free approach enabling real-time tracking of carbohydrates in live cells by using azide as a reporter under a mid-infrared photothermal (MIP) microscope (**Fig. 1a**). MIP is an emerging technique that detects the infrared (IR) absorption induced photothermal effect with a visible beam^20^. In this approach^21^, a pulsed mid-IR laser excites the chemical bond vibration. Subsequent vibrational relaxation induces a temperature rise and consequent change of local refractive index. A visible beam then detects the modulated refractive index, providing spectroscopic information of the target molecule at 300 nm resolution. Thus, by dynamic MIP mapping of azide, the trafficking of target molecules can be tracked in real time. With a commercial optical photothermal infrared microscope, mIRage, Bai *et al.* recently reported the detection of newly synthesized lipids in human-derived model systems incubated with azide-palmitic acid^22^. However, limited by the relatively low numerical aperture (NA) objective in a co-propagation geometry, the measurements were restricted to dry samples, thus lacking dynamic information.

**Fig. 1.**
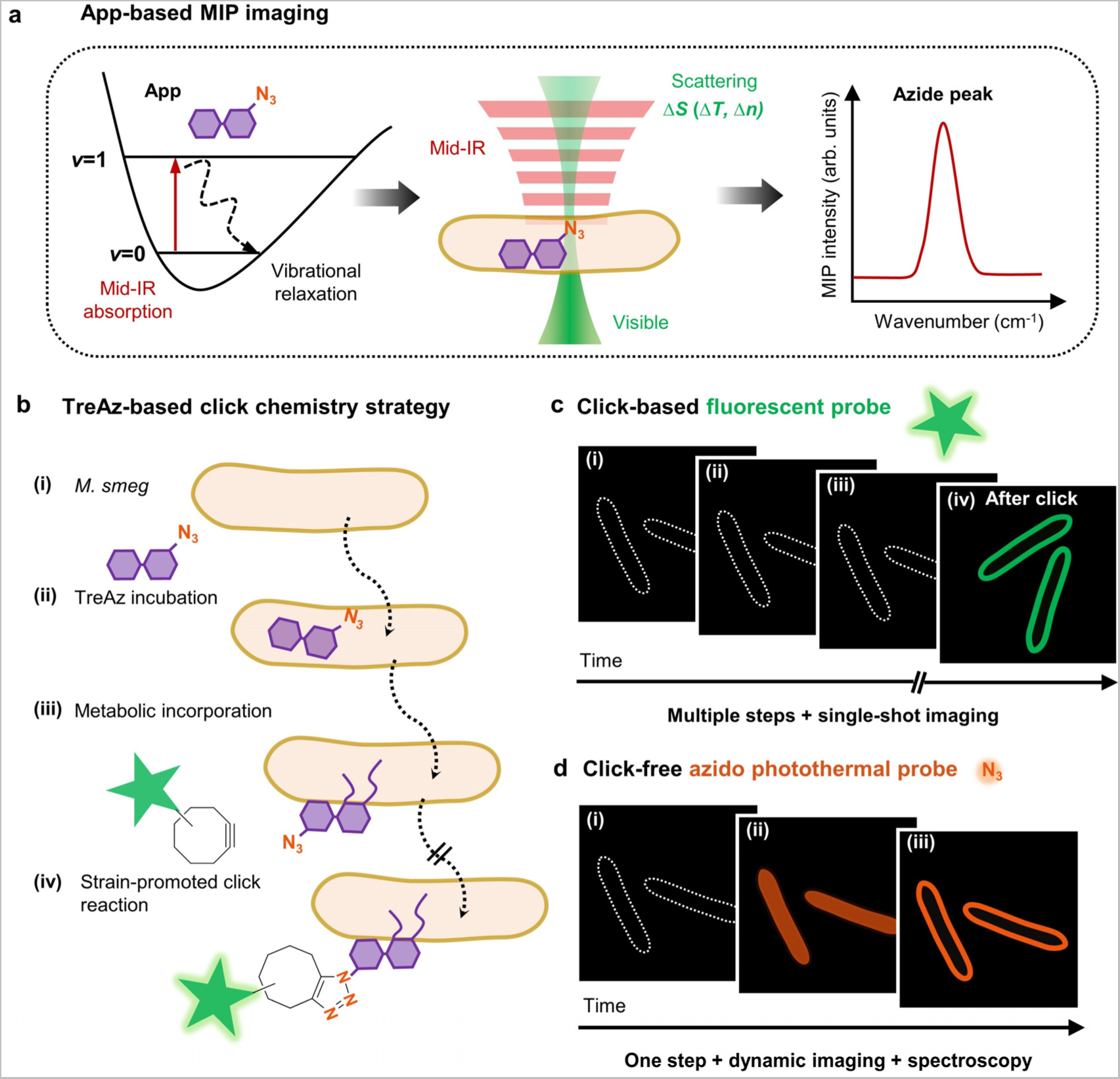
Mapping trehalose trafficking in live mycobacteria via an azido photothermal probe (App). (a) Principle of App-based mid-infrared photothermal (MIP) imaging. (b) TreAz-based click chemistry strategy for trehalose tracking in live *M. smeg*. *M. smeg*: *Mycobacterium smegmatis* mc^2^ 155. TreAz: Azido-trehalose. (c) Schematic illustration of fluorescence imaging of TreAz detection in single *M. smeg* via click chemistry. (d) Schematic illustration of click-free imaging of trehalose trafficking in live *M. smeg* via App.

We find that a counter-propagation scheme is essential to detect nanoscale carbohydrates inside a single bacterium. Moreover, we deploy a newly developed laser-scan MIP microscope^23^ for high-speed imaging of azide in live cells. We selected 6-azido-trehalose (TreAz) and *Mycobacterium smegmatis* mc^2^ 155 (*M. smeg*) as the testbed. Mycobacteria, including the pathogen *Mycobacterium tuberculosis* which is the single caustic agent to tuberculosis (TB), rely on trehalose as a crucial building block for essential cell wall glycolipids and various metabolites^11^. It is proposed that TreAz undergoes a recycling pathway and is incorporated into glycoconjugates. Previously, Bertozzi and coworkers^3^ orthogonally linked TreAz to an alkyne-functionalized fluorophore to investigate trehalose metabolic pathways in live *M. smeg* (**Fig. 1b**). However, the click reaction permanently changes the target molecule, thus only allowing a single snapshot of the molecule at the final state (**Fig. 1c**). To study specific pathways, in vitro tools were used to analyze the cell extracts^3^, but they are time-consuming.

In this work, we use TreAz as an azido photothermal probe (App) to monitor the trafficking of trehalose in mycobacteria in situ (**Fig. 1d**). Through App-based MIP imaging, we directly visualized the trehalose recycling pathway in single mycobacteria, from cytoplasmic uptake to membrane localization. By monitoring the IR peak shift of azide in MIP spectra, we show that App can be used to sense interactions between TreAz and intracellular Ag85 proteins. Intriguingly, we detected a significant amount of unreacted azide molecules after the click reaction, indicating heterogeneity of the click chemistry within a single mycobacterium and highlighting the reliability of App in tracing intracellular carbohydrates. To demonstrate the broad applicability of our approach, we further employed commercially sourced Apps to explore galactose metabolism in yeasts and glycoconjugate biosynthesis in mammalian cells. These results collectively showcase the versatility of App in unraveling complex biological processes.

## Results

### MIP spectroscopy of TreAz

To demonstrate the use of azide as a photothermal probe of carbohydrate, we performed a density functional theory calculation to quantitate the IR absorption cross section of various chemical groups conjugated to trehalose. The IR absorption cross section of azide is ∼30 times larger than nitrile and ∼90 times larger than alkyne (Supplementary Table 1). Next, we synthesized TreAz (**Fig. 2a**) as a trehalose analog due to its known metabolic pathway and its ability to replace natural trehalose^3^. Synthetic procedures of TreAz^24^ are described in Supplementary Note 1. By FTIR spectroscopy characterization of pure TreAz powder, the azide group in TreAz shows a strong IR peak at 2,105 cm^-1^ with a band width of 34 cm^-1^ (**Fig. 2b**) in the spectrally silent window (**Fig. 2c**).

**Fig. 2.**
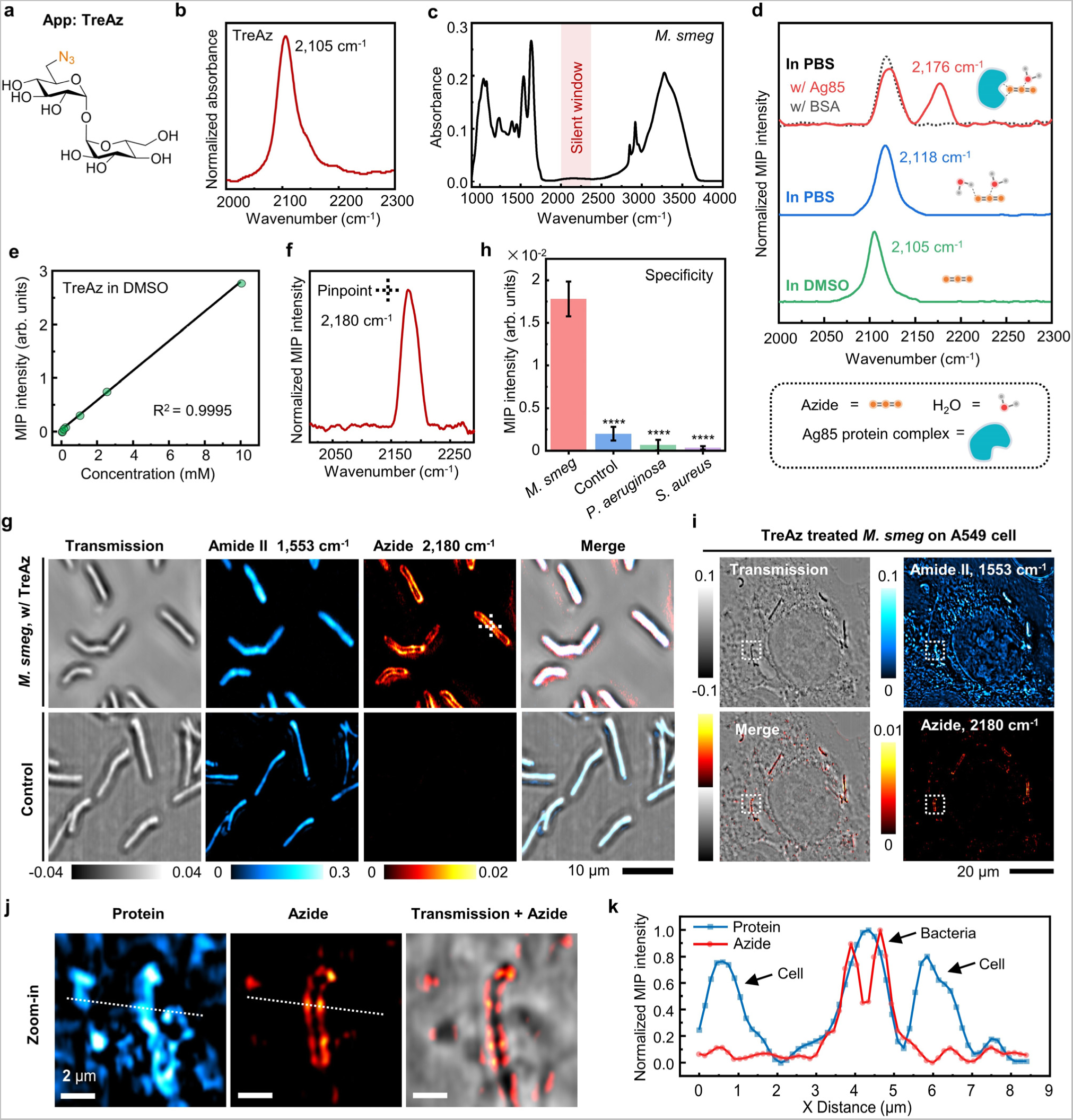
**In vitro detection and live cell MIP imaging of TreAz.** (a) Molecular structure of TreAz. Orange characters indicate the azide group. (b) FTIR spectrum of pure TreAz powder. (c) FTIR spectrum of *M. smeg*. Red region indicates the spectrally silent window, free from endogenous biomolecule peaks. (d) MIP spectra of 10 mM TreAz in DMSO (green curve), 10 mM TreAz in PBS (blue curve), 10 mM TreAz mixed with 1 mM Ag85 protein complex in PBS (red curve) and 10 mM TreAz mixed with 1 mM BSA in PBS (gray curve). (e) MIP signal intensity as a function of TreAz concentration in DMSO. (f) Pinpointed MIP spectrum of intracellular TreAz, with the pinpoint indicated by cross in panel (g). (g) MIP image of TreAz in single live *M. smeg*. Scale bar: 10 µm. (h) Specificity analysis of TreAz uptake by *M. smeg*. MIP signal intensity of TreAz treated *M. smeg* until logarithmic phase (n = 95) was compared to the signal of the control (n = 92, ****P = 8.67 × 10^-267^), *P. aeruginosa* (n = 84, ****P = 1.08 × 10^-270^) and *S. aureus* (n = 107, ****P = 4.32 × 10^-283^). Data is given as mean ± s.d. (i) Transmission and MIP image in the protein Amide II and azide channels of TreAz-treated *M. smeg* on A549 cells. Scale bar: 20 µm. (j) Zoomed-in views (white dotted boxes labeled in panel (i)) of single *M. smeg* on A549 cells. Scale bar: 2 µm. (k) MIP intensity profiles in both protein and azide channels of single *M. smeg* on A549 cells (white dotted lines labeled in panel (j)).

Additionally, the IR absorbance of azide is known to be sensitive to local environment^25, 26^. Under a co-propagating MIP microscope (Supplementary Fig. 1), the MIP spectrum of TreAz shows a distinct peak at 2,105 cm^-1^ in dimethyl sulfoxide (DMSO) (**Fig. 2d**, green curve), consistent with the FTIR spectrum. Meanwhile, a 13 cm^-1^ blue shift is observed upon changing the solvent to phosphate-buffered saline (PBS), likely due to a hydrogen-bond interaction between TreAz and water molecules^26, 27^ (**Fig. 2d**, blue curve). Notably, the Ag85 protein, an intracellular trehalose-binding protein in mycobacteria, shows strong binding with trehalose^9^. To mimic the intracellular environment, we mixed 10 mM TreAz with 1 mM Ag85 protein complex in PBS solution. The mixture of TreAz and Ag85 shows a blue-shifted peak at 2,176 cm^-1^, while the mixture of 10 mM TreAz and 1 mM bovine serum albumin (BSA) protein shows the same peak position as the TreAz in PBS at 2,118 cm^-1^ (**Fig. 2d**, red and gray curves). Thus, the large azide peak shift unveils a strong interaction between TreAz and Ag85 proteins. Such a large blueshift of the azide peak indicating the interactions between azide and proteins has also been observed by nonlinear optical spectroscopy^28, 29^. Together, the vibrational peak shift of azide highlights the potential of using App-based MIP imaging to sense intracellular molecular interactions. All MIP spectra in the silent window underwent a subtraction of solvent background as described in Supplementary Note 2 unless specially mentioned.

The MIP signal intensity of the azide is found to be linearly proportional to the TreAz concentrations with a limit of detection (LOD) around 1.3 μM in DMSO (**Fig. 2e**, Supplementary Fig. 2) and 23.6 μM in PBS (Supplementary Fig. 2). Such high detection sensitivity builds the foundation for App-based MIP imaging of carbohydrate trafficking inside live cells.

### Live cell MIP imaging of TreAz

With the high detection sensitivity, we explored counter-propagating MIP imaging of TreAz in live *M. smeg*. In this system (Supplementary Fig. 3), with a 532 nm probe beam and a high NA water immersion objective, we achieved a spatial resolution of 300 nm (Supplementary Fig. 4).

After a 26-hour incubation of 50 µM TreAz until reaching the logarithmic phase (Supplementary Fig. 5), a pinpointed MIP spectrum in the silent window (2,000 cm^-1^ to 2,300 cm^-1^) acquired inside a live bacterium shows a peak of azido groups at 2,180 cm^-1^ (**Fig. 2f**). This peak position is consistent with in vitro experiment and confirms the cellular uptake of TreAz.

For MIP imaging of intracellular azide, we chose 2,180 cm^-1^ as the on-resonance wavenumber and obtained the azide signal via subtraction of the off-resonance water background at 2,100 cm^-1^ (Supplementary Fig. 6, details in Supplementary Note 3). For TreAz-treated *M. smeg*, the azide signal is predominantly localized to the cell surface, indicating the incorporation of TreAz into the mycobacteria membrane (**Fig. 2g**). Depth-resolved MIP imaging further demonstrates TreAz localization to the membrane (Supplementary Fig. 7). For the bacteria without TreAz treatment (control group), while individual bacteria exhibit high contrast at the amide II band of 1,553 cm^-1^, indicating the distribution of proteins, no contrast at the azide channel is observed.

To test the specificity of TreAz, *Pseudomonas aeruginosa* (*P. aeruginosa*) and *Staphylococcus aureus* (*S. aureus*) were chosen as the Gram-negative and Gram-positive controls, respectively. The results show that the control, which is *M. smeg* without treating with TreAz, and the other bacterial strains treated with TreAz exhibit minimal azide signal when compared with TreAz-treated *M. smeg* (**Fig. 2h**, Supplementary Fig. 8), revealing that the uptake of TreAz is specific for *M. smeg*.

Furthermore, to demonstrate the application of App in a complex environment, we cultured the TreAz-treated *M. smeg* on A549 lung cells. As shown in **Fig. 2i** (control group shown in Supplementary Fig. 9), clear MIP contrast for single *M. smeg* on top of A549 cells were observed. The azide signal is located to the mycobacteria membrane, which is clearly distinguishable from the surrounding cell background and distinct from the protein channel (**Fig. 2j, 2k**). These results highlight that App-based MIP is capable of imaging mycobacteria in a complex environment.

Previously, Goodcare and coworkers imaged isotopically labeled single bacteria using mIRage, a commercial MIP microscope^30^. Fujita and coworkers imaged nitrile-containing molecules in single cell with a lab-built MIP microscope^31^. In both works, the single cell imaging was performed in a co-propagation manner, where the probe beam is weakly focused by a reflective objective.

We performed a head-to-head comparison between co-propagating (Supplementary Fig. 1) and counter-propagating MIP imaging (Supplementary Fig. 3) of TreAz-treated *M. smeg*. The MIP signal of single bacteria in the counter-propagating geometry shows an enhanced signal-to-noise ratio (SNR) by 11.3 times and increased resolution by 2.5 times compared to the co-propagating microscope, which allows us to resolve the membrane structure of the bacteria (Supplementary Fig. 10). Theoretically, when the focus volume for the visible beam is comparable to the thermal lens size, photothermal imaging gives the best sensitivity^32^. Thus, while co-propagating MIP is good for measuring the LOD from liquid specimens, counter-propagating MIP is the choice for high-sensitivity detection of nanoscale features in live cells.

### Visualization of trehalose recycling pathway in single mycobacteria

A recycling metabolic pathway was proposed for TreAz utilization in mycobacteria^3, 33^, as shown in **Fig. 3a**. The LpqY-SugABC transporters deliver trehalose into the cytoplasm, where trehalose is converted to trehalose monomycolate (TMM). The TMM is then transported to the periplasmic space and utilized by the protein Ag85 to synthesize trehalose dimycolate (TDM) to build the cell wall. By fluorescence microscopy, Backus *et al.* demonstrated the uptake of fluorescein-conjugated trehalose analog (FITC-Tre) by mycobacteria^9^. However, due to the large size of FITC, FITC-Tre did not enter the cytoplasm. Genetic and biochemical methods are informative but unsuitable for live cell experiments^34^. Thus, direct visualization of the trehalose metabolic pathway remains a challenge.

**Fig. 3.**
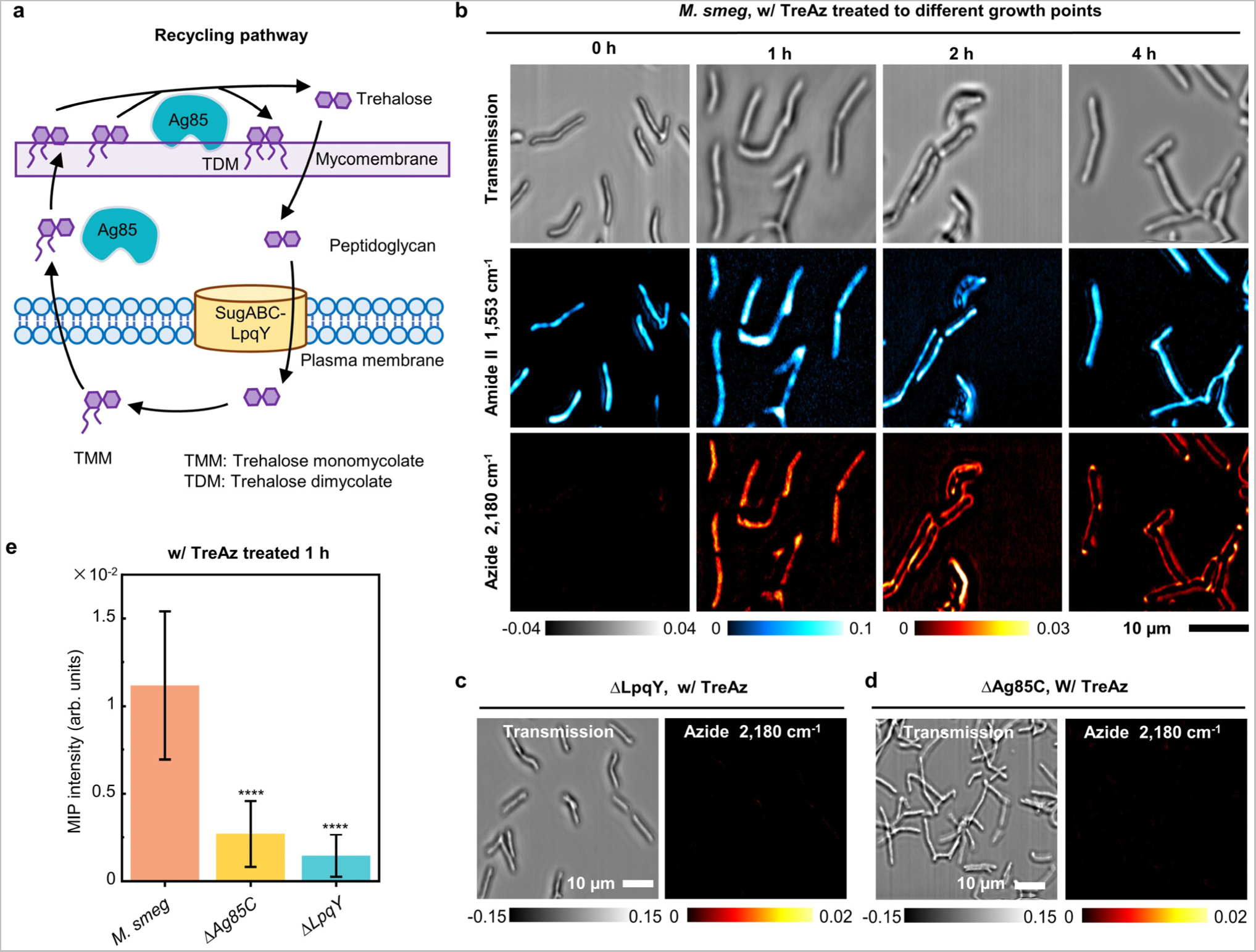
**Visualization of TreAz metabolic recycling pathway in live *M. smeg* via App-based MIP imaging.** (a) Schematics of Trehalose metabolic recycling pathway in *M. smeg*. (b) Time-course MIP imaging of live *M. smeg* treated with 50 μM TreAz in transmission mode, MIP Amide II and Azide window. Scale bar: 10 µm. (c) MIP imaging of live ΔLpqY mutant treated with 50 μM TreAz. Scale bar: 10 µm. (d) MIP imaging of live ΔAg85C mutant treated with 50 μM TreAz. Scale bar: 10 µm. (e) Quantitative azide signal of TreAz treated *M. smeg* and the mutants. MIP signal intensity of TreAz treated *M. smeg* for 1 hour (n = 89) was compared with the signal of the mutant ΔAg85C (n = 83, ****P = 8.14 × 10^-52^), and ΔLpqY (n = 77, ****P = 8.74 × 10^-60^). Data is given as mean ± s.d.

To apply the App to monitor the TreAz trafficking in mycobacteria, we performed time-course MIP imaging of logarithmic phase *M. smeg* incubated with TreAz for different time periods (**Fig. 3b**). The transmission and MIP image of proteins were recorded as references. Initially (0 hour), while the protein channel showed high contrast on the cell body, no azide signal was observed. After 1 hour of incubation, the azide signal appeared inside the cell body. By the 2-hour mark, the azide signal showed up to the cell surface. After 4 hours, the azide was predominantly localized at the poles. Such a pattern is consistent with the polar growth model of *M. smeg*, indicating active growth and division in the presence of TreAz.

Given the involvement of Ag85 and LpqY proteins in the TreAz uptake pathway^3^, we further investigated the uptake of TreAz by LpqY and Ag85C deletion mutants. The results revealed a complete abolishment of azide signals in both ΔLpqY (**Fig. 3c**) and ΔAg85C mutants (**Fig. 3d**), confirming that TreAz requires LpqY and Ag85C for entering the cytoplasm and recycling to the membrane, supported by statistical results (**Fig. 3e**). Together, App-based MIP enabled visualization of the complete TreAz recycling pathway, from uptake, membrane recycling, to pole localization.

### App-based MIP imaging allows dynamic tracking of TreAz in a single bacterium

To validate TreAz uptake, we employed click-reaction-based fluorescence imaging using a commercial fluorescent probe, AF 488 DBCO (Supplementary Note 4). This probe contains an alkyne group for the click reaction with the azido group in TreAz. *M. smeg* was treated with TreAz for 26 hours until reaching the logarithmic phase. The mycobacteria membrane was then imaged by both MIP imaging of azide before the click reaction and confocal fluorescence imaging of AF 488 DBCO after the click reaction (**Fig. 4a**). From the intensity plot profile of single *M. smeg*, clear membrane patterns were resolved by both MIP and fluorescence imaging at similar SNR (**Fig. 4b**) affirming the visualization of azide-tagged glycoconjugates on the mycobacteria membrane by App-based MIP.

**Fig. 4.**
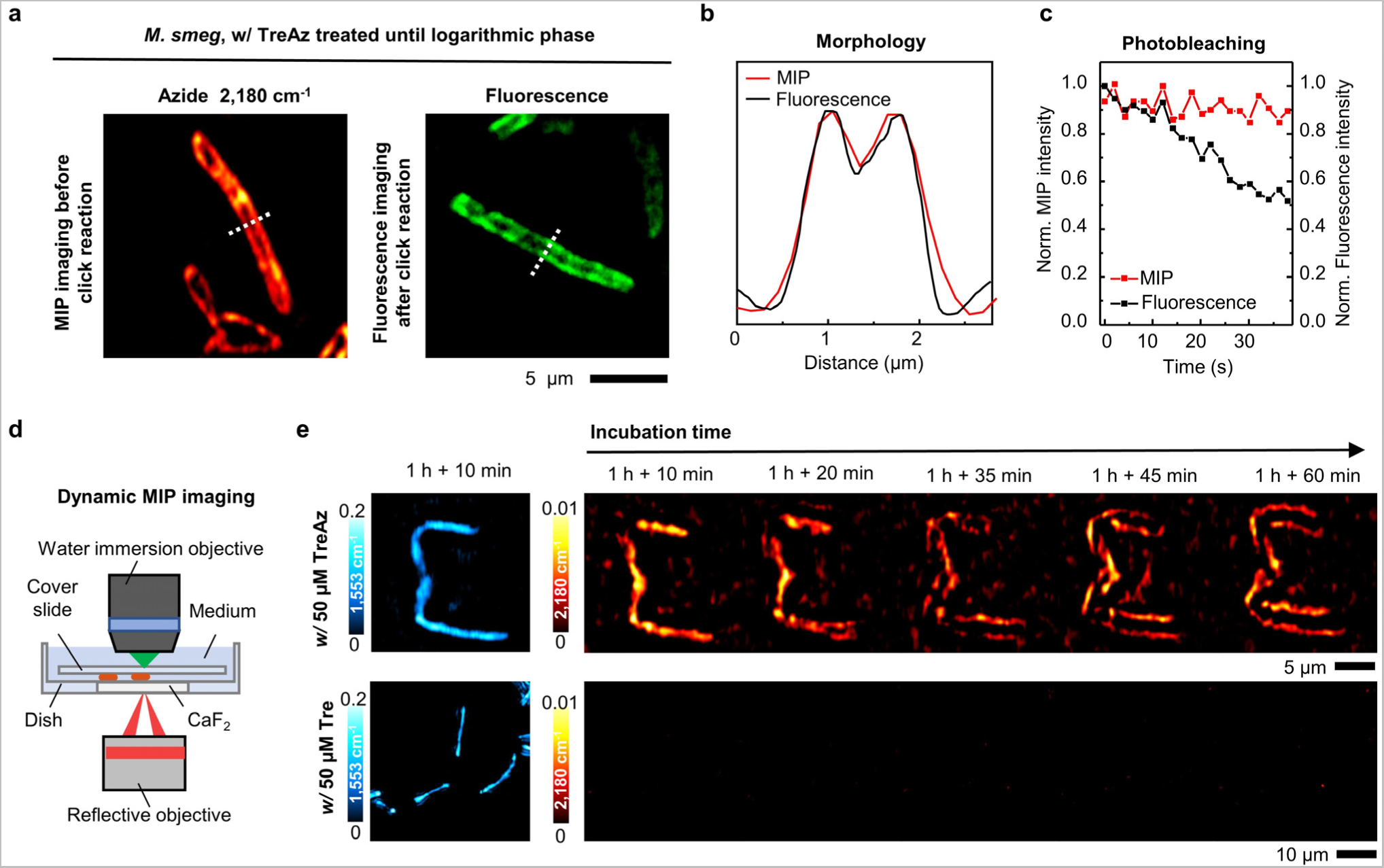
**App-based MIP imaging allows dynamic tracking of TreAz in a single bacterium.** (a) MIP imaging of 50 μM TreAz treated *M. smeg* before the click reaction and confocal fluorescence imaging of TreAz treated *M. smeg* after the click reaction. Scale bar: 5 µm. (b) Line profiles across the bacteria indicated in MIP and fluorescence imaging of *M. smeg* in (a). (c) MIP and fluorescence signal intensities versus scanning time. (d) The MIP imaging mode with a lab-made CaF_2_-bottom culture dish for longitudinal imaging of live *M. smeg*. (e) Real-time MIP imaging of TreAz trafficking in *M. smeg*. Scale bar is 5 µm in TreAz treated group and 10 µm in Tre treated group, respectively.

Notably, while fluorescence undergoes fast photobleaching, the MIP signal is resistant to photobleaching, which is essential for dynamic analysis (**Fig. 4c**). To further demonstrate the dynamic imaging capabilities of App, we made a CaF_2_-bottom culture dish suitable for longitudinal counter-propagating MIP imaging of live cells (**Fig. 4d**). Longitudinal MIP imaging of TreAz uptake into cytoplasm and recycling to membrane over 1 hour period is shown in **Fig. 4e**. Together, these results validate the trehalose recycling pathway and showcase the advantages of App-based MIP imaging.

### App-based MIP imaging reveals uneven efficiency of click chemistry in a cellular environment

As illustrated in **Fig. 5a**, the azide is converted into triazole after the strain-promoted click reaction. Thus, by MIP imaging of the remaining azide after click chemistry, one could evaluate the efficiency of the click reaction within a cell. To validate this, we added a scanning fluorescence imaging modality (Supplementary Fig. 11) to our MIP microscope, enabling multimodal fluorescence and MIP imaging of the same bacteria. Upon completion of the click reaction, the MIP signals of azido groups that undergo click reactions would disappear. Meanwhile, unabolished MIP signals indicate the unreacted azide after the click reaction.

**Fig. 5.**
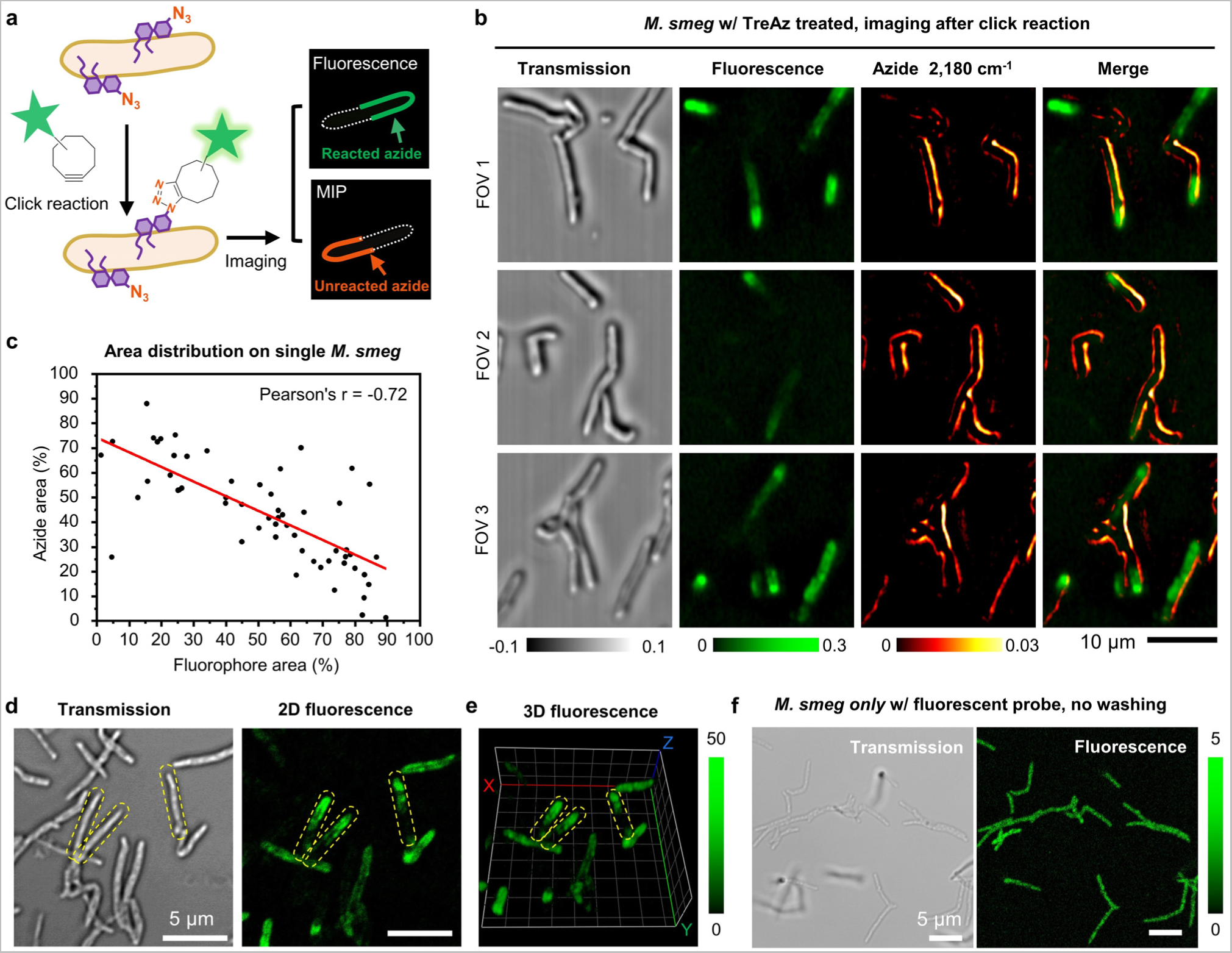
**App-based MIP imaging reveals heterogeneity of click reaction inside single *M. smeg*.** (a) Schematic illustration of complimentary MIP and fluorescence imaging contrast of TreAz treated single *M. smeg* after click reaction. (b) Colocalized MIP and fluorescence imaging of single *M. smeg* that treated with 50 μM TreAz and after click reaction. FOV: field of view. Scale bar: 10 µm. (c) Statistical analysis of azide and fluorophore area distribution on single *M. smeg* (n = 57). Pearson’s r = -0.72, indicating statistically significant negative correlation. (d) Uneven fluorescence distribution was observed in a commercial confocal fluorescence imaging of single *M. smeg* treated with TreAz and underwent click reaction. Yellow dotted lines indicate single bacteria. Scale bars: 5 µm. (e) Depth-resolved confocal fluorescence imaging of TreAz-treated *M. smeg* after click reaction in panel (d). Yellow dotted lines indicate single bacteria. Each grid represents a size of 2 µm. (f) Confocal fluorescence imaging of *M. smeg* only with fluorescent probe incubation. Scale bars: 5 µm.

We incubated *M. smeg* with TreAz until the logarithmic phase and followed a standard protocol of click chemistry^3^. Strikingly, we observed a significant amount of MIP signal from azide after the click reaction in different fields of view (**Fig. 5b**) and two independent repeats (Supplementary Fig. 12). Statistically, a significantly negative correlation between fluorescence signals and MIP signals of azide was observed on individual bacteria (**Fig. 5c**, Pearson’s r = -0.72). To explore whether the unreacted azido groups truly come from the heterogeneity of click reaction, a series of validation experiments were conducted. Firstly, we repeated the same experiment with a commercial confocal fluorescence microscope. A similar pattern of uneven fluorescence distribution was found (**Fig. 5d**). Next, depth-resolved confocal fluorescence imaging of *M. smeg* after click reaction was performed. In 3D confocal fluorescence, a portion of the cells displayed uneven fluorescence signals along the cell body (**Fig. 5e**), eliminating the influence of non-uniformity caused by different focal planes. Finally, to evaluate the impact from the diffusion of fluorescent probes, *M. smeg* were incubated only with the fluorescent probe and imaged without washing. Uniform fluorescence intensity was observed along individual bacteria, indicating good permeability and uniform diffusion of the fluorescent probe inside the bacteria (**Fig. 5f**). Collectively, these data reveal the heterogeneity of strain-promoted click chemistry in the mycobacteria membrane environment.

The observed heterogeneity in the click reaction within mycobacteria could potentially be attributed to the relatively large size of dibenzocycloctynes in the clickable dye AF 488 DBCO. It has been reported that the dibenzocycloctynes could not be site-specifically incorporated into proteins^35^, and the sterically encumbered nature of the molecule may limit the effectiveness of the click reaction in bioconjugation applications^36^. To further explore the origin of the heterogeneity, logarithmic phase *M. smeg* was incubated with TreAz until different growth points, and then analyzed either via MIP imaging of the azide, or by fluorescence imaging of the fluorescent probe after click reaction (Supplementary Fig. 13). After 0, 1, 2 and 4-hour incubation of TreAz, MIP imaging demonstrated a high resolution to differentiate the locations of the azide reporter at different time points. In comparison, the fluorescence images displayed a weak but uniform contrast at 1 hour and uneven intensity along individual bacteria at 2 hours and 4 hours. These results indicate click reaction is more favorable in the cytoplasm and on the poles.

### App-based MIP imaging of carbohydrates in various biological systems

Benefiting from the rapid development of click chemistry, a variety of azide-tagged carbohydrate products are commercially available or with well-defined synthetic procedures^37, 38^. To demonstrate the versatility of Apps in other biological systems, we explored galactose metabolism in yeast *Saccharomyces cerevisiae* (*S. cerevisiae*) and glycoconjugate biosynthesis in mammalian HeLa cells by harnessing commercially sourced azido-carbohydrates.

Galactose serves as a vital carbon source for energy production in yeast cells, primarily metabolized through the Leloir pathway^39^. The pathway initiates with the conversion of β-D-galactose (β-D-Gal) into α-D-galactose (α-D-Gal) catalyzed by a mutarotase enzyme, followed by phosphorylation of α-D-Gal. The enzymatic phosphorylation of galactose into galactose-1-phosphate (Gal-1-P) exhibits a preference for the α-form of the galactose^40^, where α and β anomers of galactose differ in the stereochemistry of the C-1 carbon atom (**Fig. 6a**).

**Fig. 6.**
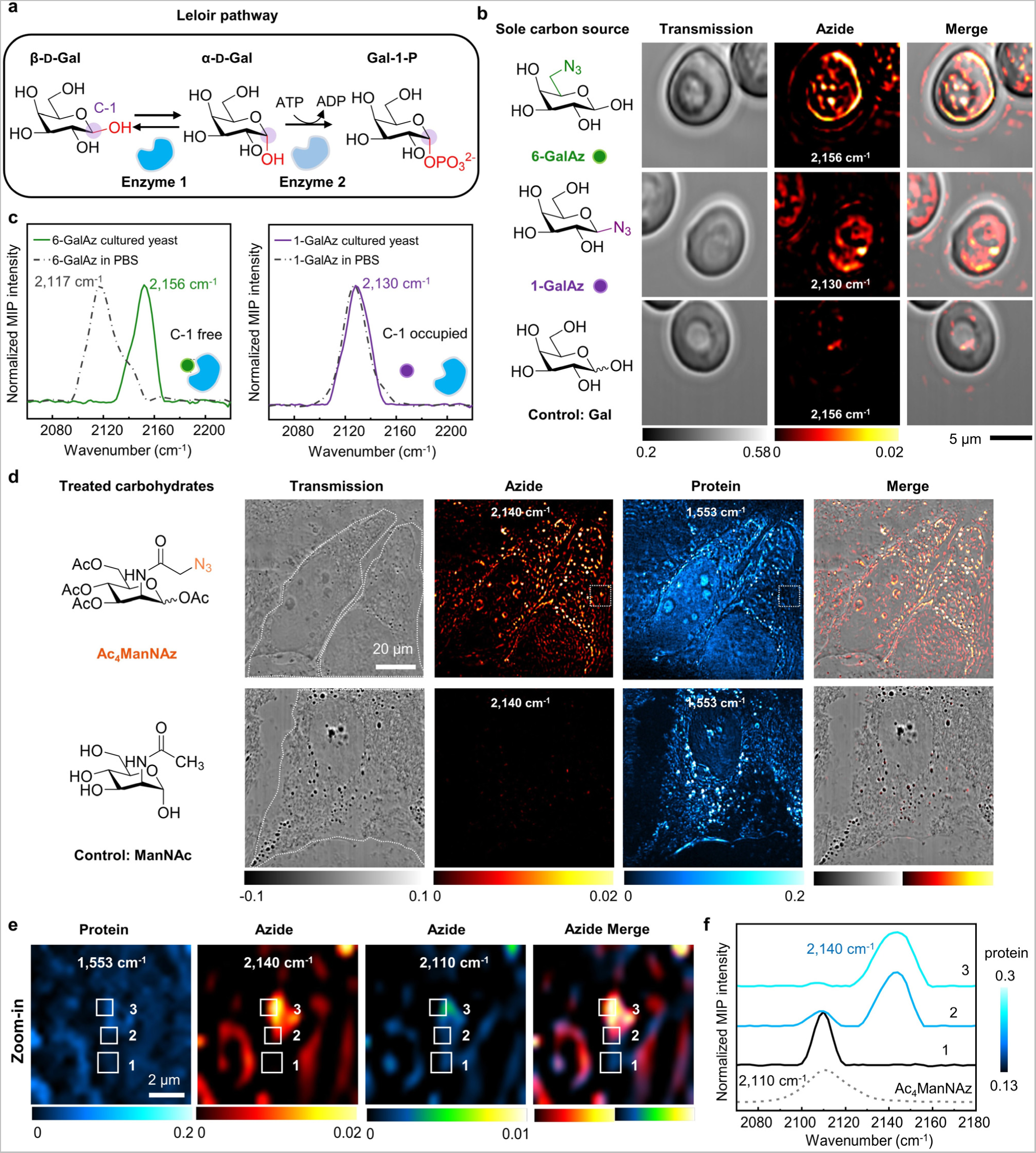
**App-based MIP detection of carbohydrate metabolism and interaction with proteins in various biological systems.** (a) Initial steps of Leloir pathway for galactose metabolism in yeast. β-D-Gal is reversibly converted into α-D-Gal via mutarotase (Enzyme 1), where the α and β anomers of galactose differ in the stereochemistry of the C-1 carbon atom, labeled with purple circles. α-D-Gal is then phosphorylated to Gal-1-phosphate (Gal-1-P) via galactokinase (Enzyme 2), which can subsequently be converted into a derivative of glucose to generate energy. (b) MIP imaging of live yeast cells cultured with 2% 6-GalAz, 1-GalAz, and Gal for 24 hours. Scale bar: 5 µm. The experiment was repeated independently three times with similar results. (c) Validation of interactions between proteins and galactose in Leloir pathway via azide peak shift. Solid curves are MIP spectra acquired from GalAz cultured yeast cells. Dash-dotted curves are MIP spectra of 50 mM GalAz in PBS solution. (d) MIP imaging of 50 μM Ac_4_ManNAz and ManNAc treated HeLa cells in azide and protein channels. Scale bar: 20 µm. White dotted lines in transmitted images indicate the outline of the cells. The experiment was repeated independently three times with similar results. (e) Multicolor MIP imaging of Ac_4_ManNAz treated HeLa cell in protein, azide and merge channels. The imaging areas are zoomed-in views of white dotted boxes labeled in panel (d). (f) Spectroscopic correlation between azide peak shift and protein abundance in Ac_4_ManNAz treated HeLa cell. The MIP spectra and protein profile were acquired from the white boxes labeled area in panel (e). Gray dotted line indicates the MIP spectrum of 10 mM Ac_4_ManNAz in ethanol.

To visualize the Leloir pathway in live cells, we employed two azide-tagged galactoses: 6-azido-6-deoxy-D-galactose (6-GalAz) and 1-deoxy-β-D-galactopyranosyl azide (1-GalAz), with the azide at C-6 and C-1, respectively (**Fig. 6b**). Yeast cells cultured in medium with 6-GalAz showed a sharp azide peak at 2,156 cm^-1^, while the cells cultured with 1-GalAz presented a clear azide peak at 2,130 cm^-1^

(Supplementary Fig. 14). Live-cell MIP imaging revealed a concentrated distribution of 6-GalAz and a diffuse distribution of 1-GalAz (**Fig. 6b**). Control cells cultured with D-galactose (Gal) without azide conjugation showed no peaks in the silent region. Fluorescence imaging of the yeast cells after click reactions further substantiated the MIP results, confirming the uptake of 6-GalAz and 1-GalAz (Supplementary Fig. 15).

Moving beyond imaging, we explored protein-galactose interactions in the Leloir pathway through azide peak shifts (**Fig. 6c**). An obvious azide peak shift of 6-GalAz from 2,117 cm^-1^ in PBS to 2,156 cm^-1^ in the yeast cell is observed, indicative of carbohydrate-enzyme binding^41^. In contrast, 1-GalAz showed no azide peak shift between solution and live cell MIP spectrum, indicating that the azide group occupying the C-1 position hinders its further metabolism in the Leloir pathway. Consequently, 1-GalAz exhibited a diffuse pattern inside the cell. These findings provide visual evidence of the interactions between carbohydrates and proteins during the Leloir pathway. Together, MIP detection of peak shift in the azide vibration offers a new way to study intracellular carbohydrate-enzyme interaction, which is beyond the reach of click-reaction-based fluorescence imaging.

We further extended App-based MIP imaging to monitor glycan synthesis in mammalian cells. For glycan visualization, tetraacylated N-azidoacetylmannosamine (Ac_4_ManNAz) has been widely used for metabolic labeling of glycoconjugates in cells and mice through click reaction^15, 42, 43^. To demonstrate the applicability of App to mammalian cells, we performed MIP imaging of Ac_4_ManNAz to map glycosylation in HeLa cells (**Fig. 6d**). *N*-Acetyl-D-mannosamine (ManNAc) was used as a control. HeLa cells treated with 50 µM Ac_4_ManNAz presented a strong azide peak at 2,140 cm^-1^ in the MIP spectrum, whereas the control cells showed no peak in the silent region (Supplementary Fig. 16). In Ac_4_ManNAz-treated cells, the azide signals were predominantly localized to the cell surface and inside the cells, as confirmed by their positions in the transmitted images (**Fig. 6d**, Supplementary Fig. 17). No azide contrast shows up in the control group. The MIP images at 1,553 cm^-1^ provides a reference map of protein distribution inside cells. Our result reveals Ac_4_ManNAz incorporation into the newly synthesized azido-glycoconjugates inside HeLa cells and on the cell surface, consistent with the glycoconjugate biosynthesis process^44^. Click-reaction-based fluorescence imaging validated the MIP results (Supplementary Fig. 18).

Importantly, the high spatial resolution and high sensitivity of App-based MIP imaging allowed observation of two distinct peaks of azide-tagged carbohydrates across a single protein-rich component (**Fig. 6e**). As the MIP intensity of proteins increased in the selected areas labeled as the position from 1 to 3 (**Fig. 6e**), the azide peak at 2,110 cm^-1^ attenuates, and a shifted azide peak at 2,140 cm^-1^ shows up (**Fig. 6f**). As a reference, the MIP spectrum of 10 mM Ac_4_ManNAz in ethanol exhibited a peak at 2,110 cm^-1^. Together, this spectroscopic data suggests a transition from free states to protein-bound states of the azido-carbohydrate molecules.

## Discussion

Click chemistry-based fluorescent imaging has become a powerful method for studying carbohydrate metabolisms. In this work, we introduce App-based MIP imaging that enables dynamic mapping of cellular entry, intracellular trafficking, and metabolism of carbohydrates without the need for click chemistry. App-based MIP imaging of intracellular carbohydrates in various biological systems, from single bacteria, yeasts, to mammalian cells, are demonstrated. Trehalose recycling in single mycobacteria, from cytoplasmic uptake, membrane recycling, to pole localization, was directly visualized. Strikingly, we observed a significant amount of unreacted azide after following a standard protocol of click reaction, indicating click chemistry heterogeneity and incompleteness within a single mycobacterium. Moreover, through monitoring the shift of the azide vibrational peak in the silent window, we show that App can be used to sense intracellular carbohydrate-protein interactions, which is beyond the reach of fluorescence imaging.

We note that click-free vibrational imaging was firstly demonstrated via Raman imaging of EdU. By using the alkyne-tagged cell proliferation probe, Yamakoshi *et al.* detected newly synthesized DNA in HeLa cells. Yet, a long acquisition time of 49 minutes for one image (∼50 by 50 µm) were needed^45^. Stimulated Raman scattering (SRS) microscopy much improved the imaging speed and offered a LOD of 200 µM for alkyne-tagged molecules^46^. Compared to Raman scattering, IR absorption benefits from a much larger cross section^21^. Wei and coworkers reported direct IR imaging of azides in single cells, yet with low spatial resolution of around 3 µm^47, 48^. MIP microscopy offers both high resolution and high sensitivity through the use of a tightly focused visible probe beam. To compare MIP and SRS imaging of azide, we performed SRS spectroscopic detection of pure TreAz powder, which exhibited a Raman peak of the azide group at 2,105 cm^-1^. However, for 2.5 mM TreAz dissolved in DMSO, no SRS peak of azide vibration was distinguished from the DMSO background (Supplementary Fig. 19). We further performed SRS imaging of *M. smeg* treated with 50 µM TreAz for 26 hours. Although SRS showed a high contrast at the C-H channel, the azide signals in single bacterium were not detectable by SRS microscopy (Supplementary Fig. 19). These results demonstrate the sensitivity advantage of App-based MIP imaging, which is essential for mapping low-concentration biomolecules in a living system.

One limitation of App-based MIP imaging lies in the water photothermal background in the silent window. The water molecules exhibit a weak absorption centered around 2100 cm^-1^, contributed by the water bend-libration combination band^49^. In this work, we used the difference between on- and off-resonance images to map the azide in biological cells. This method requires the acquisition of two MIP images upon tuning the quantum cascade laser. A more elegant way of distinguishing MIP signals from the water photothermal background is to use thermal dynamics^50^. Compared to nanoscale objects, water exhibits a much larger thermal decay constant. Thus, by signal digitization^50^ or two-window boxcar detection^51^, time-resolved measurement of thermal dynamics can be pursued to extract the signal from water background upon excitation with a single IR pulse per pixel.

For future work, as TreAz works well for MIP imaging of trehalose uptake by mycobacteria with a high specificity, it can be potentially used for detecting individual mycobacteria in clinical samples, aiding in rapid diagnosis of TB infection. Another important direction will be real-time MIP imaging of glycan synthesis in vivo upon treatment of live tissues or animals with azido-carbohydrates. The concept of App can be extended to other molecules other than carbohydrates. For instance, azido-amino acids can be used to facilitate the study of protein synthesis and function in live cells. We envision App-based MIP imaging will pave a new way for investigation of the localization, trafficking, and metabolism of biomolecules in life and diseases.

## Methods

### Cell lines and materials

*M. smeg* mc^2^ 155 and the mutants were obtained from the Biodefense and Emerging Infections Research Resources Repository (BEI Resources). *Pseudomonas aeruginosa* (*P. aeruginosa*) 1133, *Staphylococcus aureus* (*S. aureus*) ATCC6538, A549 cell line and HeLa cell line were obtained from ATCC. *S. cerevisiae* strain (W303) was obtained from a published strain^52^. Ag85 Complex (NR-14855) was obtained from BEI Resources. Bovine serum albumin (BSA, A7030), dimethyl sulfoxide (DMSO, 472301), D-(+)-Trehalose dihydrate (Tre, 90210), Luria-Bertani broth (LB, 1102850500) and yeast nitrogen base (YNB, 51483) were purchased from Sigma Aldrich. Ethanol (A962-4) was purchased from Fisher Scientific. Complete supplement mixture (CSM, 1001-010) was obtained from Sunrise Science Products. Middlebrook 7H9 (271310) and ADC enrichment (212352) were purchased from BD Bioscience. Glycerol (AC158922500) was purchased from Acros Organics. AF 488 DBCO (218F0) was purchased from Lumiprobe Corporation. 6-GalAz (AF432) and 1-GalAz (GL631) were purchased from Synthose Inc. Gal (48260-100G-F), Ac_4_ManNAz (900917-50MG) and ManNAc (A8176-250MG) were purchased from Sigma Aldrich. TreAz was synthesized and purified according to a previous procedure^24, 53^. CaF_2_ substrates (ø 25 mm and 1mm thick, CAFP25-1) were purchased from Crystran.

### MIP microscopy

For LOD measurement, an upright co-propagating MIP microscope was used (Supplementary Figure 1). The system was constructed on an inverted microscope frame (IX73, Olympus), integrating a continuous-wave 532 nm laser (Samba, HUBNER photonics) for the visible probe and a pulsed quantum cascade laser (MIRcat 2400, Daylight Solutions) tunable from 900 cm^-1^ to 2,300 cm^-1^ as the mid-IR pump. The visible probe was co-aligned with the mid-IR pump laser and focused into a sample by a reflective objective lens (40X, 0.5NA, LMM440x-P01, Thorlabs). The probe photons were collected in forward direction. Two photodiodes (PD, DET100A2, Thorlabs) were used to acquire the transmission and MIP images, respectively. For MIP detection, the current from the photodiode was converted to voltage through a 50 Ω resistor (T4119, Thorlabs), and filtered by a high pass filter (0.12-1000 MHz, ZFHP-0R12-S+, Mini-Circuits), then amplified by two low noise amplifiers (gain = 40 dB, 1 kHz-500 MHz, SA-251F6, NF corporation). Finally, the signal was filtered by a low pass filter (BLP-1.9+, DC-1.9 MHz, 50 Ω, Mini Circuits), then delivered to a lock-in amplifier (HF2LI, Zurich) to demodulate the MIP signals.

For intracellular imaging, a counter-propagating beam geometry on the same upright MIP microscope was used (Supplementary Figure 3). The IR beam was focused on the sample with a reflective objective lens (40X, 0.5NA, LMM440x-P01, Thorlabs) and the visible beam was focused by a water immersion objective lens (60X, 1.2 NA, UPlanSApo, Olympus). A galvanometer mirror (Saturn 1B, ScannerMax) was used to scan the visible beam. Simultaneously, the IR beam was synchronously scanned by another pair of galvanometer mirrors (GVS001, Thorlabs). The probe photons were collected in the forward direction using a visible/IR dichroic mirror (GEBBAR-3-5-25, Andover Corporation), and the intensity was sensed by a photodiode (PD, DET100A2, Thorlabs). For longitudinal MIP imaging of App within the same cells, an inverted counter-propagating MIP microscope was utilized. In this setup, a lab-built frame was used. The IR beam was directed from the bottom, while the visible light was focused from the top. The accessories were the same as those used in the upright counter-propagating MIP microscope.

For LOD measurement, the IR laser was set with a repetition rate of 100 kHz and pulse width of 100 ns to reduce heat accumulation during long-time spectral scanning. The mid-IR laser continuously swept wavenumbers from 2,000 cm^-1^ to 2,300 cm^-1^ at a speed of 50 cm^-1^/s. The raw MIP spectra were obtained from a lock-in amplifier with a time constant of 20 ms. For MIP imaging of bacteria, the IR laser was set with a repetition rate of 1 MHz, a pulse width of 80 ns, and an average power after the objective ranging from 8-20 mW, depending on the wavenumber. An average power of the probe laser was ∼50 mW after the objective. The MIP images were acquired with a pixel dwell time of 10 µs and a step size of 150 nm. For hyperspectral MIP imaging of yeast and HeLa cells, the IR laser was set with a repetition rate of 290 kHz and pulse width of 100 ns to reduce heat accumulation during long-time scanning. The hyperspectral scanning range was from 2,060 cm^-1^ to 2,220 cm^-1^ with a step size of 2 cm^-1^/frame. The MIP images were acquired with a pixel dwell time of 25 µs and a step size of 200 nm (yeast) and 300 nm (HeLa cell), respectively.

### LOD

The LOD determines the minimum analyte concentration at which the MIP signal can be reliably distinguished from the background noise, and is defined as:

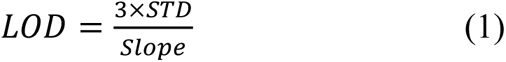

where *STD* is the standard deviation of the MIP intensity in the blank solvent solution, and *Slope* is the slope of the calibration curve relating concentration to MIP signal. To get the calibration curve, MIP spectra of TreAz were obtained at various concentrations in PBS and DMSO solutions, respectively. A 2 µL solution was sandwiched between two CaF_2_ substrates, and the MIP spectra were acquired via a co-propagating MIP system. This geometry ensured that the focus volume of the probe beam was comparable to the thermal lens of the solution medium, providing an ideal MIP signal for LOD detection^32^. After background subtraction of the solvent spectra and baseline correction, the MIP intensity at azide peak was used to establish the calibration curve between concentration and MIP signal. The slope values were determined as 0.0168 mM^-1^ for PBS and 0.277 mM^-1^ for DMSO measurements. The *STD* was measured from the MIP intensity of the solvent at the corresponding azide peak (2,118 cm^-1^ for PBS and 2,015 cm^-1^ for DMSO). Specifically, *3 x STD_PBS_ = 3.97 x 10^-4^* and *3 x STD_DMSO_ = 3.74 x 10^-4^*. According to Equation 1, the LOD of TreAz is determined to be 1.3 μM in DMSO and 23.6 μM in PBS.

### FTIR measurement

The FTIR spectra of all samples were measured on an attenuated total reflection FTIR spectrometer (Nicolet Nexus 670, Thermo Fisher Scientific). The spectra resolution is 2 cm^-1^ and each spectrum was measured with 128 scanning. All spectra were automatically normalized by the baseline correction on the system. Additionally, for samples in solvent, the background spectra of the correspoding solvent were subtracted.

### Fluorescence imaging

The fluorescence imaging was pursued by a laser-scanning fluorescence modality integrated into the MIP microscope. An additional 488 nm laser (06-MLD, Cobolt) was co-aligned with the 532 nm laser to serve as the excitation source, with the laser power output set at 12 mW. The power of the excitation laser on sample was around 0.1 mW. A 60x water immersion objective with NA of 1.2 was used (UPlanApo, Olympus) as MIP imaging. The fluorescence emission was collected in an epi-direction with a filter set (Excitation filter: FES0500, Thorlabs; Dichroic beam splitter: Di03-R405/488/532/635-t1-25×36, Sermock; Emission filter: FF01-525/30-25, Sermock).

The emitted photons were collected by a photomultiplier tube (PMT, H10721-110, Hamamatsu). The transmitted images were collected using the MIP imaging mode. The confocal fluorescence images were acquired on a Zeiss LSM 880 laser scanning microscope equipped with a ×63/1.4 NA oil immersion objective and ZEN software was used to collect the data.

## Data processing

All images were analyzed and displayed with ImageJ. The data were plotted using Origin. Pseudocolor was added to the MIP and fluorescent images with ImageJ. Individual bacterium was marked for the quantification of single-cell MIP signals, except for the measurement presented in **Fig. 5b**, where the MIP and fluorescence intensities on the cell surface were measured. The transmitted images were normalized by the subtraction of the background. 3D reconstruction was done with ImageJ using the 3D viewer plugin. The detailed MIP spectra and images processing workflow are shown in Supplementary Note 2 and Supplementary Note 3, respectively.

### Bacteria preparation

Middlebrook 7H9 broth supplemented with ADC enrichment was prepared using Middlebrook 7H9 (4.7 g), glycerol (2.0 mL), and distilled water (900 mL). The mixture was then autoclaved (Tuttnauer EZ 10, Hauppauge, NY) for sterilization before use. Following autoclaving, ADC enrichment (100 mL) was added to the sterilized Middlebrook 7H9 broth once its temperature reached 45 °C. *M. smeg* was cultured in 7H9 liquid medium in incubation tubes at 37 °C with shaking until the optical densities measured at wavelength of 600 nm (OD600) reached 0.5, typical within 22-26 hours (Supplementary Fig. 5). A stock solution of 2.5 mM TreAz in PBS buffer was prepared. To 1.96 mL of *M. smeg* in an incubation tube, 40 μL of TreAz stock solution was added, which gave the TreAz concentration of 50 μM. Subsequently, *M. smeg* was incubated at 37 °C for different incubation time of 0-hour, 1 hour, 2 hours and 4 hours while shaking at 250 rpm. For the control group, *M. smeg* was treated with Tre with the same procedure.

To culture *M. smeg* on A549 lung cells, A549 cells were seeded on CaF_2_ substrates in a culture dish with a density of 1×10^5^ CFU mL^-1^ with 2 mL culture medium overnight at 37°C and 5% CO_2_. After cell attachment, cells were washed with PBS three times, then fixed with 4% formaldehyde. The cells were finally washed with PBS for two times. *M. smeg* cells was added into the dish with 7H9 liquid medium and 50 μM TreAz, and incubated with the A549 cells for 2 hours at 37°C. For the control group, no TreAz was added to the culture dish. Finally, the samples were sandwiched between a CaF_2_ substrate and a cover glass for imaging.

*P. aeruginosa* and *S. aureus* were cultured in LB medium at 37 °C with shaking until OD600 reached 0.5. To 1.96 mL of *P. aeruginosa* and *S. aureus* in incubation tube, 40 μL of TreAz stock solution was added, which gave the TreAz concentration of 50 μM. The bacterial cells were then incubated at 37 °C for 1 hour while shaking at 250 rpm.

After collection, the bacteria were centrifuged (3500 rpm, 3 minutes) and washed three times with PBS. About 1-2 µL of concentrated bacteria were sandwiched between a CaF_2_ substrate and a cover glass ready for imaging. For dynamic MIP imaging, about 10 µL of concentrated bacteria were added to a lab-made CaF_2_-bottom culture dish containing 200 µL PBS with 50 μM TreAz or Tre. To prevent bacterial movement, a glass cover slide was added to the center of the dish.

### Yeast cell preparation

The synthetic defined (SD) medium was prepared by mixing 79 mg CSM, 746 mg YNB and 100 mL distilled water, and was sterilized by autoclaving before use. Yeast cells were grown 3 days in SD medium with with 2% (w/v) ethanol at 30 ℃ with shaking without other carbohydrates. After collection, the cells were centrifuged (3000 rpm, 5 minutes) and washed twice with sterilized distilled water. Then, the cells were incubated in SD medium with 2% (w/v) Gal or GalAz as the sole carbon source for one day at 30 ℃ with shaking. After collection, the cells were centrifuged (3000 rpm, 5 minutes) and washed three times with sterilized distilled water. About 1 to 2 µL of concentrated yeast cells were sandwiched between the special CaF_2_ substrates and the cover glass for imaging.

### HeLa cell preparation

HeLa cells were seeded on CaF_2_ substrates with a density of 1×10^5^ ml^-1^ with 2 mL minimum essential medium overnight at 37°C and 5% CO_2_. After cell attachment, the cells were treated with Ac_4_ManNAz or ManNAc by adding 1.25 μL (for a final concentration of 50 μM Ac_4_ManNAz or ManNAc) of 10 mM Ac_4_ManNAz or ManNAc in ethanol stock solution to each dish. Cells were incubated with Ac_4_ManNAz or ManNAc for 40 hours at 37°C. Cells were then washed with PBS three times, then fixed with 4% formaldehyde. The cells were finally washed with PBS for two times and sandwiched under a piece of cover glass for imaging.

### Click chemistry

The click reaction was performed following an established protocol^3^. 200 μL of bacterial cells or yeast cells were transferred to a 0.5-mL sterile centrifuge tube, followed by centrifugation (3500 rpm, 3 minutes, 4 °C) and triple washing with PBS with 0.5% BSA (PBSB). Subsequently, the cells were incubated with AF 488 DBCO (Lumiprobe, Catalog: 218F0) (1:1000 dilution of 1 mM stock solution in DMSO into PBSB) for one hour at room temperature in the dark. After centrifugation (3500 rpm, 3 minutes, 4 °C) and washing with PBSB for three times, the cells were fixed with 200 μL 4% paraformaldehyde in PBS in the dark for 10 minutes. Final washing with PBS two times readied the cells for analysis by fluorescence or MIP microscopy.

For click reaction in HeLa cells, after treatment of Ac_4_ManNAz or ManNAc, the cells were rinsed with live cell imaging solution (A14291DJ, Invitrogen) for two times. Each dish was filled with 50 μM AF 488 DBCO (diluted from a 10 mM DMSO stock solution) in solution, and incubated for 1 hour at 37 °C in the dark. The cells were washed for three times with PBS and fixed the cells with 4% formaldehyde for 15 minutes at room temperature in the dark. The cells were finally washed with PBS for two times and sandwiched under a piece of cover glass ready for imaging.

### Statistics & reproducibility

All experiments were independently repeated at least three times with similar results. The sample sizes for all statistical experiments exceeded 7 per group. All data collected during the experiments were included. No data were excluded from the analyses. Statistical analysis was performed using Origin version 2018. All groups were expressed as mean ± s.d. Column means were compared using one-way ANOVA. When ANOVA showed a significant difference, pair wise comparisons between group means were examined by Bonferroni comparison test. Significance was defined when p < 0.05.

## Data availability

The data supporting the findings of this study are available within the article and supplementary information. Source data are provided with this paper.

## Acknowledgments

This research was supported by NIH grants R35GM136223, R01AI141439, and R33 CA261726 to JXC. The authors thank Giulio Chiesa for providing the yeast strain. The authors thank Meng Zhang for helpful discussions on the bacteria culture and Jianpeng Ao for helpful discussions on the manuscript and SRS imaging.

## Author contributions

Q.X., M.Y. and J.-X.C. conceived the concept and designed the experiments. Q.X. performed the experiments and analyzed the data. R.B. cultured the HeLa cell and performed the cell treatment. H.A.P. synthesized TreAz and helped with *M. smeg* culture. J.Y. developed the counter-propagating scanning MIP system. Z.G. helped with the scanning fluorescence imaging system. M.L. helped with the co-propagating scanning MIP system. Z.G., J.Y. and X.G. coded the program for data acquisition. H.H. helped with the confocal fluorescence imaging. Z.P. and Q.C. did the calculation of IR cross section. X.G. performed the SRS imaging. O.R. intellectually contributed to the experiment design. Q.X. and J.-X.C. co-wrote the manuscript. All authors have read and approved the manuscript.

