## SUPPLEMENTARY INFORMATION for "Click-free imaging of carbohydrate trafficking in live cells using an azido photothermal probe"

Summary

Number of Pages: 25

Pages 2-3: Supplementary Notes 1-4
Pages 4: Supplementary Scheme 1
Page 5: Supplementary Table 1
Pages 6-24: Supplementary Figs. 1-19
Page 25: Supplementary References

**Supplementary Note 1. Synthetic procedures of TreAz.**

The synthetic procedure of 6-azido-trehalose (TreAz) is shown as follows:

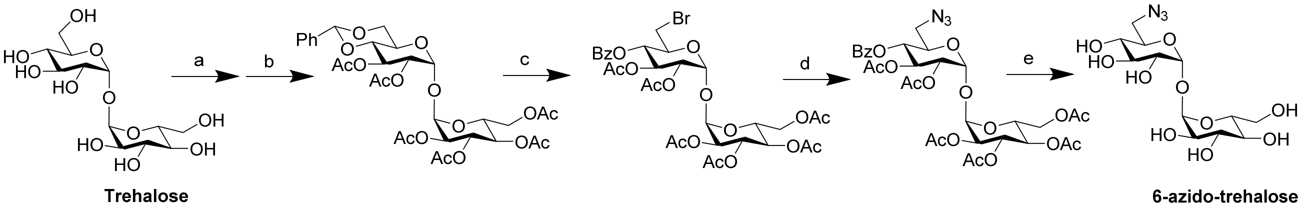


(a) $PhCH(OCH_3)_2$, PTS, DMF, 25 °C for 18 h and 40 °C for 4 h; (b) $Et_3N$, $Ac_2O$, DMAP, DMF, 25 °C, overnight (40% overall yield from combined steps of a and b); (c) NBS, $CaCO_3$, trifluorotoluene, 77 °C, 5 hours (80%); (d) $NaN_3$, DMF, Ar, 65 °C, 5 hours (77%); (e) $NaOCH_3$, $CH_3OH$, 25 °C, 1.5 hours (97%). The detailed characterizations of product and intermediates are reported in previous work[1].

**Supplementary Note 2. MIP spectra acquisition and processing workflow.**

The MIP spectra for both solutions and bacteria were obtained from the pinpoint MIP spectra using previous method[2]. To acquire pinpoint MIP spectra of solutions, the focused visible beam was centered in the solution based on the transmitted image, and the mid-IR laser continuously swept wavenumbers from 2,000 cm$^{-1}$ to 2,300 cm$^{-1}$ at a speed of 50 cm$^{-1}$/s. The raw MIP spectra were obtained from a lock-in amplifier with a time constant of 20 ms. Since both DMSO and water have the absorbance in this region, all MIP spectra in the cell-silent window underwent a subtraction of solvent background or background from the control samples. The spectra then further underwent the baseline correction and smoothing with the Savitzky-Golay method using Origin. The final MIP spectra were normalized by the IR power profile in the corresponding region. For pinpoint MIP spectra of bacteria, both the cell surface and cell cytosol were selected as target points of interest based on the transmitted images, and control spectra were acquired from untreated bacteria. The workflow is shown as follows:

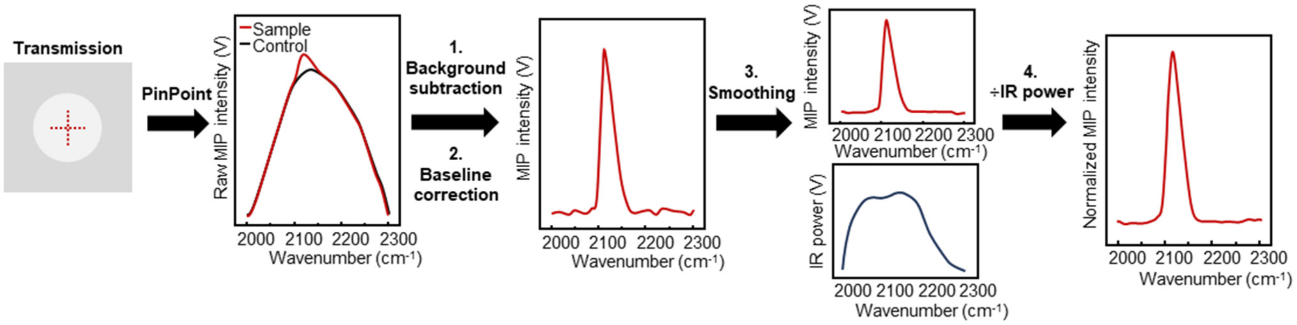


To investigate the biological systems with large sizes, it is hard to select the area for acquiring pinpoint MIP spectra of App treated samples. Thus, all the MIP spectra for yeast and HeLa cells were obtained from the hyperspectral MIP images. The hyperspectral scanning range is from 2,060 cm$^{-1}$ to 2,220 cm$^{-1}$ with a step size of 2 cm$^{-1}$/frame. The intracellular MIP spectra were then obtained from the selected region of interest (typically 25 pixels) in the hyperspectral MIP imaging stack. The final MIP spectra were obtained by background subtraction, baseline correction, smoothing and normalized by the IR power profile in the corresponding region. The workflow is shown as follows:

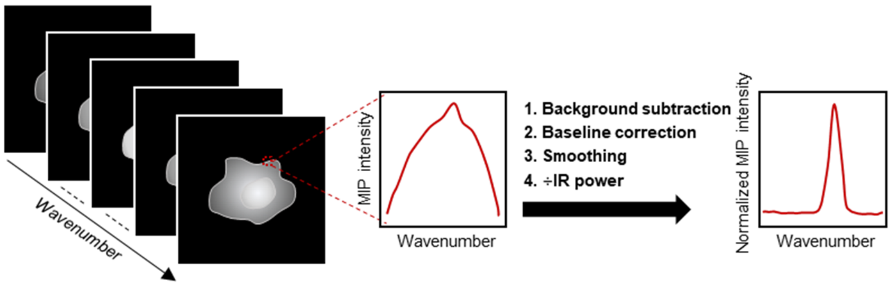


**Supplementary Note 3. MIP image processing method.**

To obtain the MIP images of App treated bacteria at cell silent window, water background subtraction was performed using previous method[2]. To remove the water background, two selected frames at the azide peak of TreAz at 2,180 $cm^{-1}$ and the water infrared absorbance at 2,100 $cm^{-1}$ were collected, respectively. The wavenumbers of 2,100 $cm^{-1}$ and 2,180 $cm^{-1}$ were selected based on the MIP spectra of TreAz treated bacteria and control bacteria from both of the cell surface and cytosol (Supplementary Fig. 6). The azide treated bacteria show an obvious peak at 2,180 $cm^{-1}$ above the background absorption. The MIP signal intensity in the control sample at 2,100 $cm^{-1}$ is equal to the MIP signal intensity at azide peak of 2,180 $cm^{-1}$ in the same control sample. After subtraction, the bacteria in the control groups exhibit no signal (Fig. 2g). Thus, all MIP images of bacteria were collected at 2,180 $cm^{-1}$ and underwent a subtraction process using MIP images collected at 2100 $cm^{-1}$ to remove the water background and to generate the final MIP images. To enhance signal-to-noise ratio, the images at all channels were obtained via the averaging of 10 frames.

For MIP imaging of yeast and HeLa cells, the hyperspectral MIP images were denoised using a BM4D V3.2 (http://www.cs.tut.fi/~foi/GCF-BM3D/index.html#ref_software) denoising algorithm[3, 4] via MATLAB. Then, water background subtraction was applied to the single-color MIP image based on the corresponding MIP spectra of the yeast cells (Supplementary Fig. 14). For 6-Gal treated and control yeast, MIP images in the azide channel were generated by subtracting the MIP image at 2,156 $cm^{-1}$ (azide peak) from the half sum of the MIP images at 2,130 $cm^{-1}$ and 2,170 $cm^{-1}$ (water absorbance). For 1-Gal treated yeast, MIP images in the azide channel were generated by subtracting the MIP image at 2,130 $cm^{-1}$ (azide peak) from the half sum of the MIP images at 2,110 $cm^{-1}$ and 2,150 $cm^{-1}$ (water absorbance). For azide treated and control HeLa cells, MIP images in the azide channel were generated by subtracting the MIP image at 2,140 $cm^{-1}$ (azide peak) from the half sum of the MIP images at 2,120 $cm^{-1}$ and 2,160 $cm^{-1}$ (water absorbance) according to the MIP spectra of the cells (Supplementary Fig. 16).

**Supplementary Note 4. Click-reaction-based fluorescence imaging.**

The clickable fluorescent dye used in this work is a commercial product **AF 488 DBCO**. The alkyne group in dibenzocyclooctyne (DBCO) can react with the azide group in azide-tagged molecules, such as TreAz, and form a stable triazole via strain promoted alkyne-azide cycloaddition[5]. The scheme is illustrated as follows:

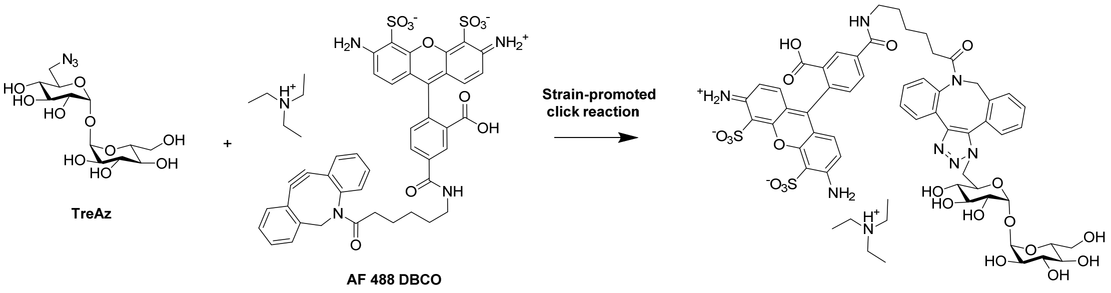

### Supplementary Scheme 1. Molecular structures.

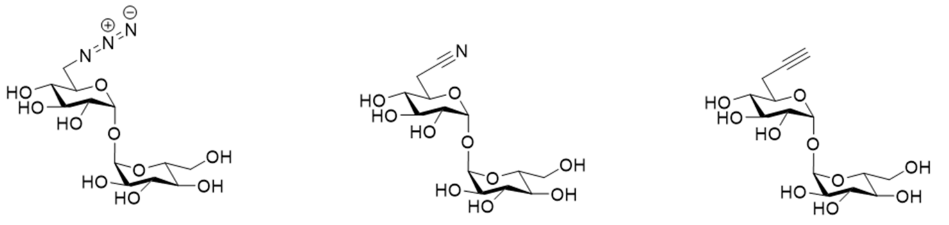

Azide tagged trehalose (TreAz)    Nitrile tagged trehalose (TreCN)    Alkyne tagged trehalose (TreCC)

**Supplementary Table 1. IR absorption peak cross sections of azide, nitrile and alkyne conjugated to trehalose.** Calculations were performed using the B3LYP/6-311G** level of theory[6, 7, 8] with the Gaussian 16 Revision C.01 Program. Initial structures were built with Avogadro Version 1.2.0[9]. These structures were then optimized by Gaussian, prior to a frequency calculation. Molecular structures are shown in Supplementary Scheme 1.

| Trehalose analogs | IR Cross section ($10^{-17}$ $cm^2$/molecule) |
|---|---|
| TreAz | 2.61 |
| TreCN | 0.079 |
| TreCC | 0.029 |

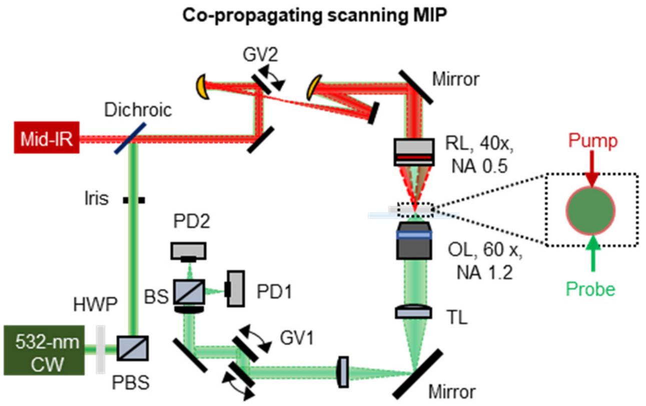


**Supplementary Fig. 1 Schematic illustration of the co-propagating scanning MIP imaging system.** The visible probe was co-aligned with the pump mid-IR laser and focused by a reflective objective (RL). Probe photons were collected in forward direction. The intensity was sensed by a silicon photodiode (PD). PD1 was used to acquire the transmission and PD2 was used to acquire the MIP image. BS: T90:R10 beam splitter, GV1 and GV2: galvo mirrors, HWP: half-wave plate, NA: numerical aperture, OL, objective lens, PBS: polarized beam splitter, TL: tube lens. Inside the dotted box is the schematic illustration of the beam size of pump and probe.

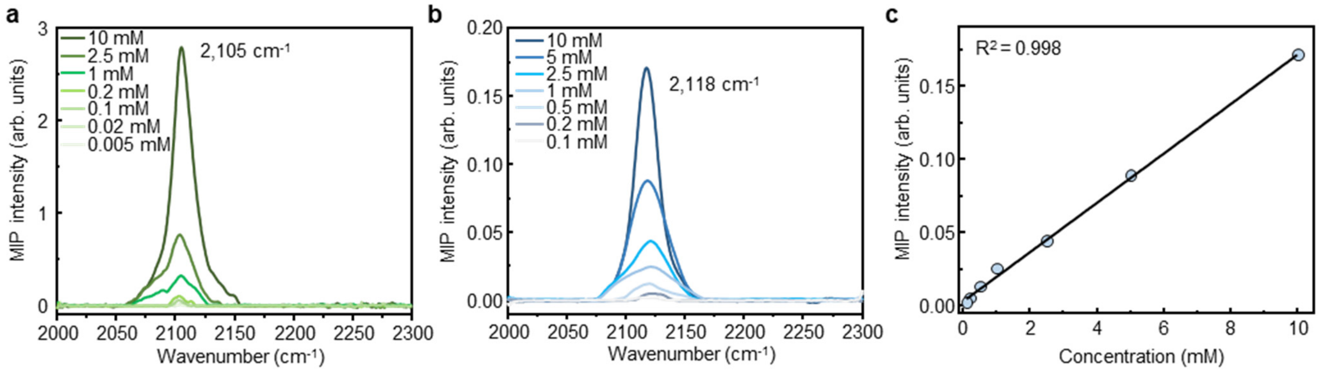


**Supplementary Fig. 2 Limit of detection (LOD) of TreAz.** (a) MIP spectra of TreAz at different concentrations in DMSO. (b) MIP spectra of TreAz at different concentrations in PBS. (c) MIP signal intensity of TreAz at different concentrations in PBS.

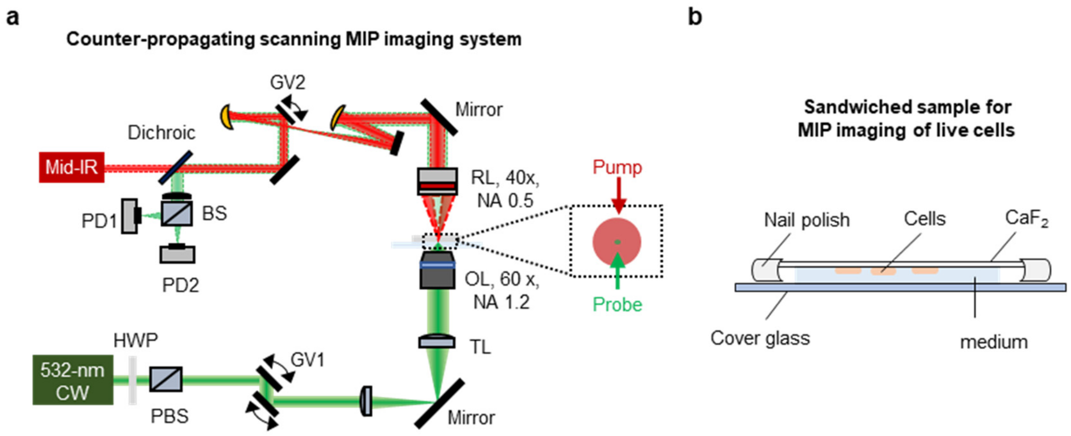


**Supplementary Fig. 3 Schematic illustration of counter-propagating scanning MIP imaging system.** (a) MIP microscope setup. The IR pump beam was generated by a tunable (from 900 to 2,300 $cm^{-1}$) quantum cascade laser. The visible probe was provided with a continuous-wave 532 nm laser. Probe photons were collected in forward direction. The intensity was sensed by a silicon photodiode (PD). PD1 was used to acquire the transmission and PD2 was used to acquire the MIP image. BS: T90:R10 beam splitter, GV1 and GV2: galvo mirrors, HWP: half-wave plate, NA: numerical aperture, OL, objective lens, PBS: polarized beam splitter, RL: reflective objective, TL: tube lens. Inside the dotted box is the schematic illustration of the beam size of pump and probe. (b) Illustration of sandwiched sample for MIP imaging of live cells. Top substrate is $CaF_2$, and the bottom substrate is No. 1 glass cover slide. Cells were sandwiched between two substrates and immersed in medium.

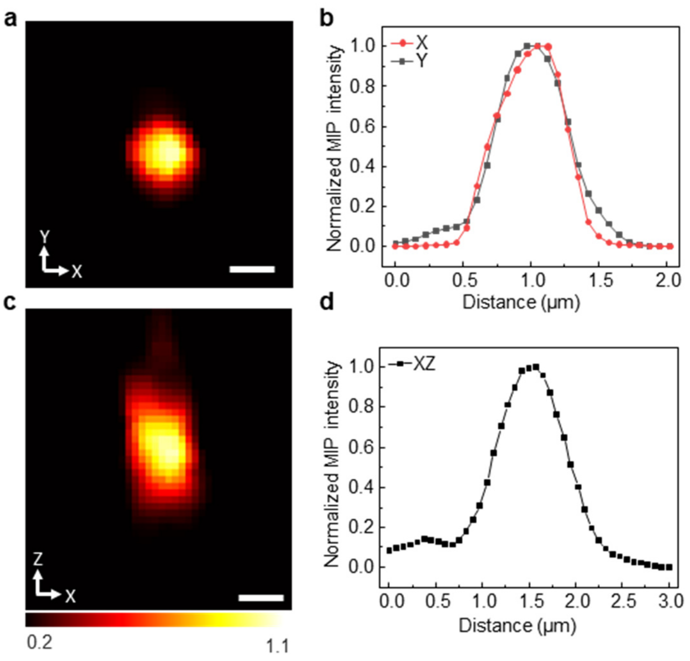


**Supplementary Fig. 4 Spatial resolution of the counter-propagating MIP imaging System.** (a) MIP image of a 500-nm diameter PMMA particle acquired with IR excitation at 1,729 $cm^{-1}$. (b) Line profiles of MIP intensity along X and Y axes, with fitted full width at half maximum (FWHM) of 578 nm (X) and 584 nm (Y), respectively. (c) Projection of the same particle along the XZ axis. (d) Line profiles of MIP intensity along XZ axis, with a fitted FWHM of 823 nm. The 3D point spread function is deconvolved using the image with the actual particle size along the three principal axes, resulting in a lateral resolution of 290 nm (X) and 302 nm (Y), and an axial resolution of 653 nm. Scale bars: 500 nm.

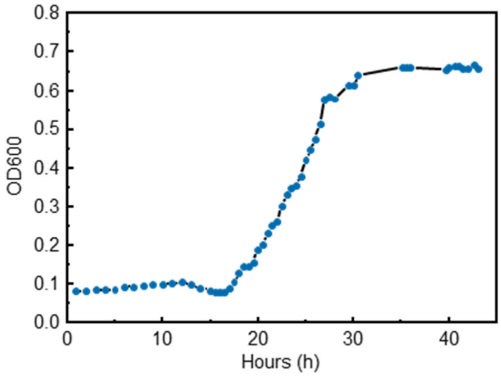


**Supplementary Fig. 5 A representative growth curve of *M. smeg*.** OD600 represents optical densities measured at 600 nm wavelength.

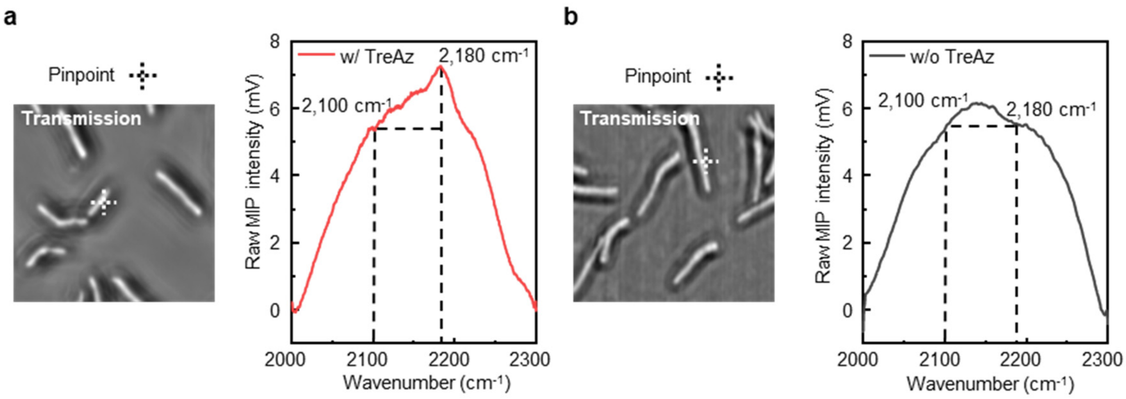


**Supplementary Fig. 6 Raw pinpoint MIP spectra of *M. smeg* in the silent region.** MIP spectra acquired from (a) TreAz treated *M. smeg* and (b) control *M. smeg*. The spectra shown here are raw data without water background subtraction.

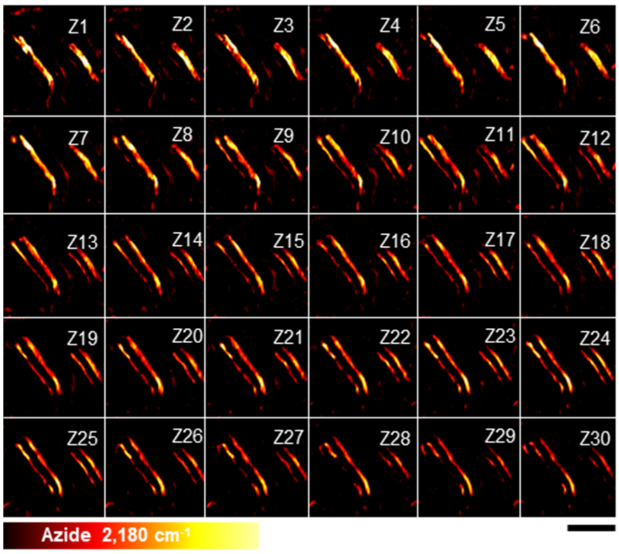


**Supplementary Fig. 7 MIP Z stack imaging demonstrates TreAz localization to the membrane.** Multi-stack MIP imaging of TreAz treated *M. smeg* at the azide channel. The Z1 image indicates the bottom of the bacteria. The Z step size is 100 nm. Scale bar: 10 µm.

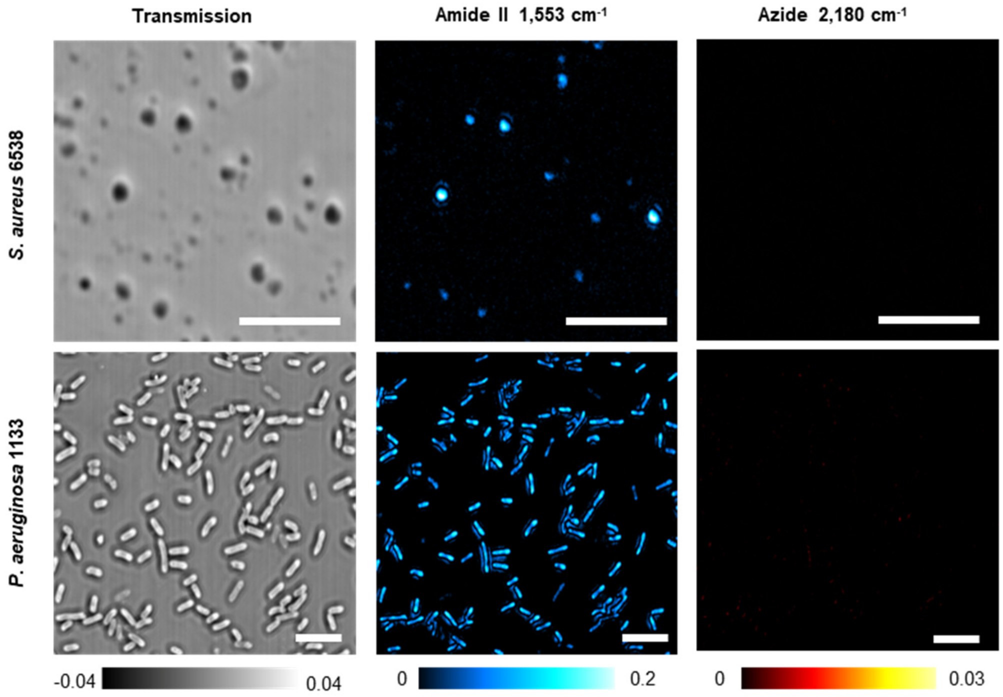


**Supplementary Fig. 8 MIP imaging of control bacterial strains treated with TreAz.** Gram-positive *S. aureus* 6538 and Gram-negative *P. aeruginosa* 1133 were both treated with 50 µM TreAz for 1 hour prior to MIP imaging. Scale bars: 10 µm.

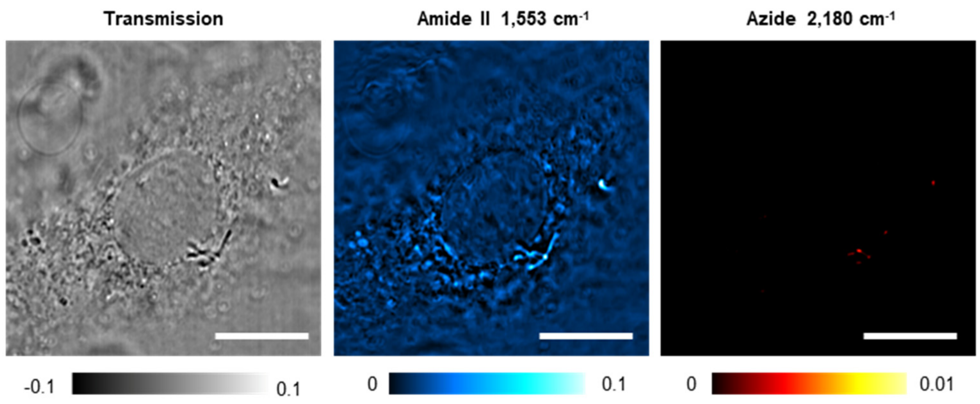


**Supplementary Fig. 9 MIP imaging of untreated *M. smeg* deposited on lung A549 cells.** Scale bars: 20 μm.

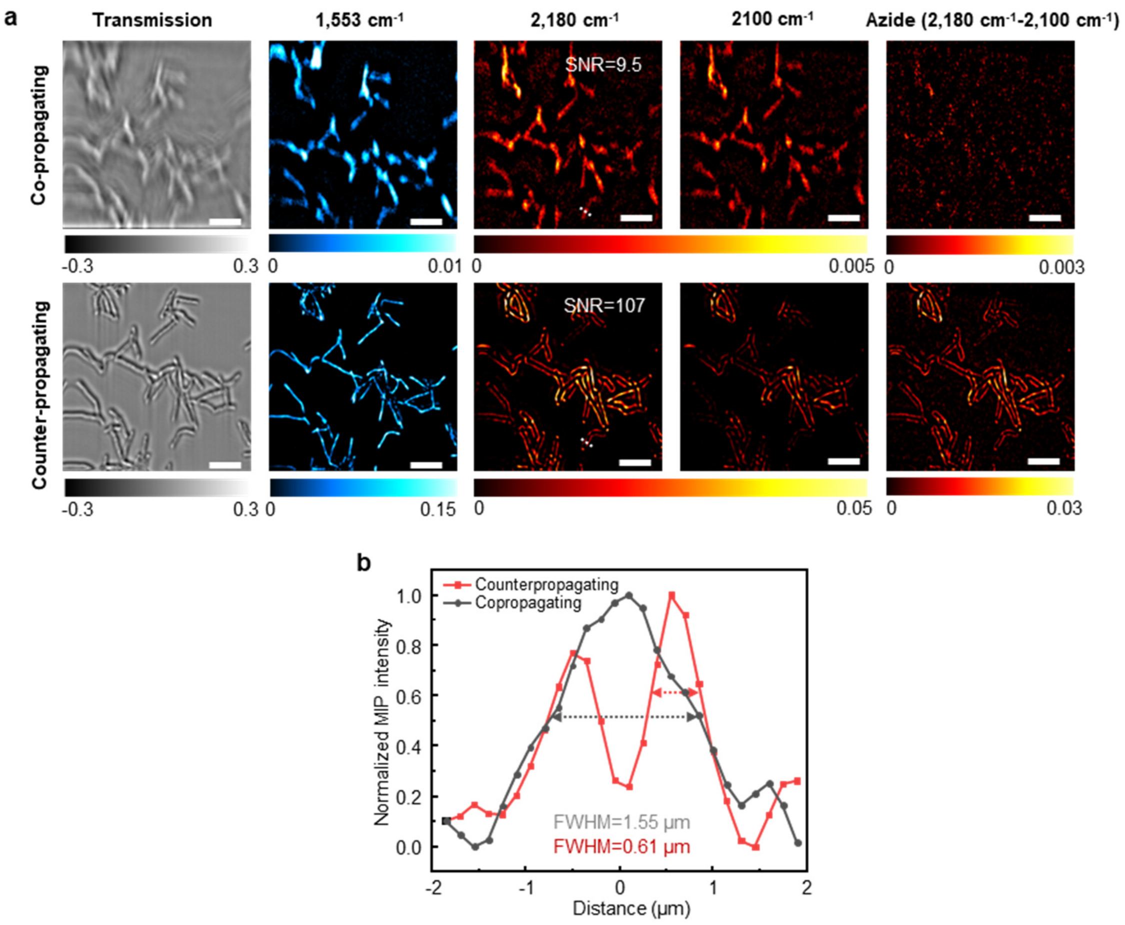


**Supplementary Fig. 10 Head-to-head comparison between counter-propagating and co-propagating MIP imaging of TreAz treated *M. smeg*.** (a) Multicolor MIP imaging of TreAz treated *M. smeg*. Treatment time: 26 h. The signal-to-noise ratio (SNR) level at 2180 $cm^{-1}$ was calculated from the ratio between the mean value of selected cell region (4 pixels) and the standard deviation of the selected background region (4 pixels) in single color MIP imaging at 2,180 $cm^{-1}$. Scale bars: 5 µm. The MIP images at 1553 $cm^{-1}$ shows protein distribution. After subtracting the off-resonance image at 2110 $cm^{-1}$ from the on-resonance image at 2180 $cm^{-1}$, the azide signal from the mycobacterial membranes is buried in the noise in the co-propagating system but is clearly seen in the counter-propagating system, showing the significance of using a tightly focused probe beam to sense nanoscale objects in MIP microscopy. (b) Line profiles of MIP intensity along the cell body labeled with dotted white lines in panel a (2180 $cm^{-1}$). The full width at half maximum (FWHM) is 1.55 µm in co-propagating imaging and 0.61 µm in counter-propagating, respectively.

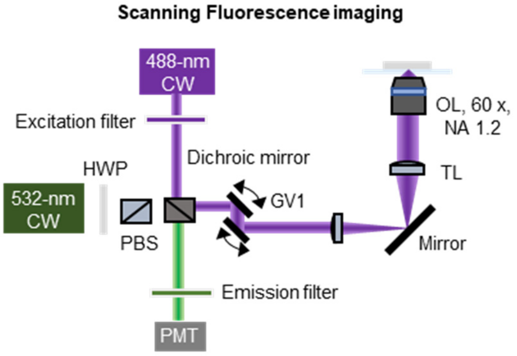


**Supplementary Fig. 11 The lab-built fluorescence imaging modality on the MIP microscope.** The excitation light was provided with a continuous-wave 488 nm laser. Emission was collected in epi-detection via a photomultiplier tube (PMT). The transmitted image was collected with the illumination of 532 nm laser via the same pathway for MIP detection in Supplementary Fig. 3. GV1: galvo mirrors, HWP: half-wave plate, NA: numerical aperture, OL: objective lens, PBS: polarized beam splitter, TL: tube lens.

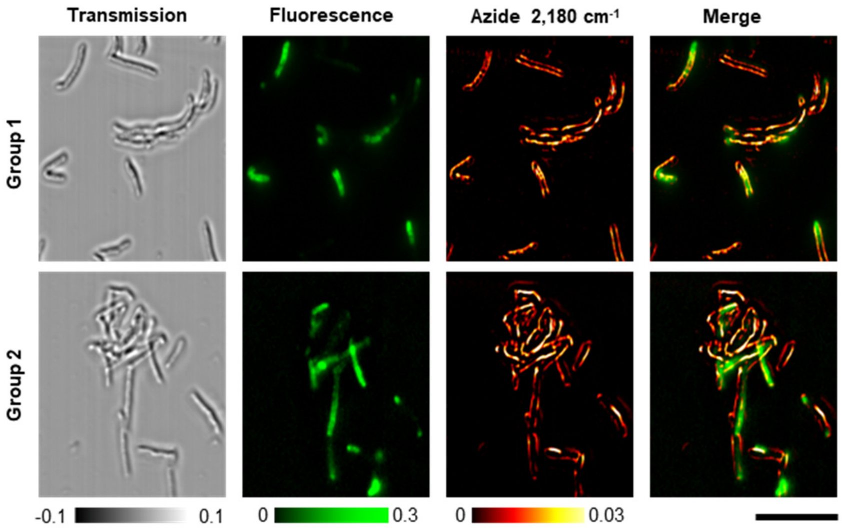


**Supplementary Fig. 12 Two independent repeats of colocalized MIP and fluorescence imaging of single *M. smeg*.** The bacteria were treated with 50 µM TreAz till the logarithmic phase, then underwent click reaction following a standard protocol of click chemistry. Scale bar: 20 µm.

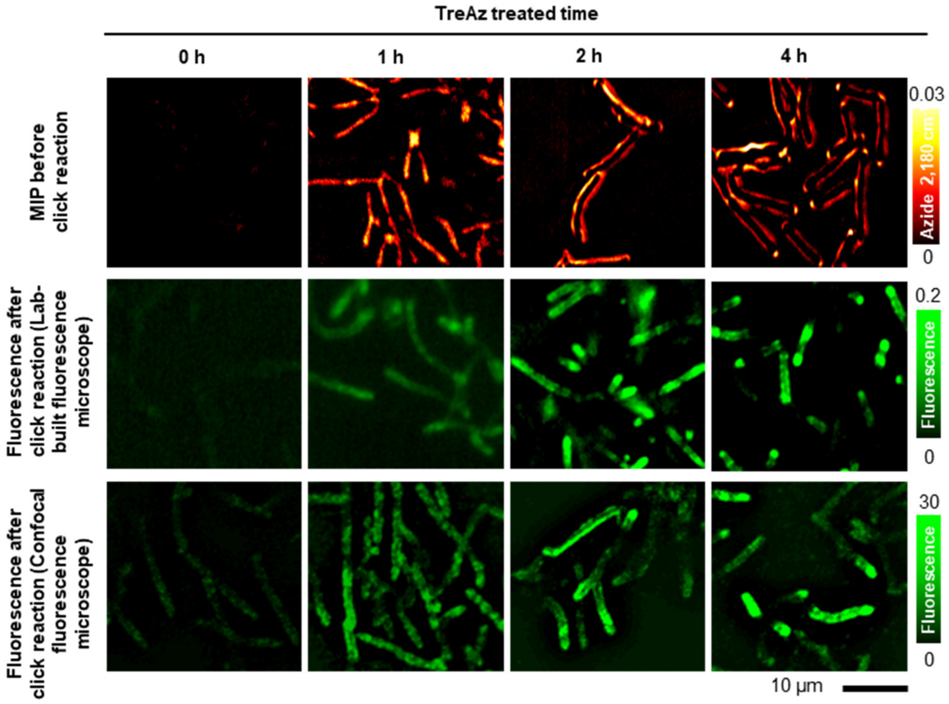


**Supplementary Fig. 13 MIP and fluorescence imaging of TreAz treated *M. smeg* with different incubation times.** Fluorescent mages were acquired on a lab-built fluorescence microscope and a commercial confocal fluorescence microscope, respectively. Scale bar, 10 µm.

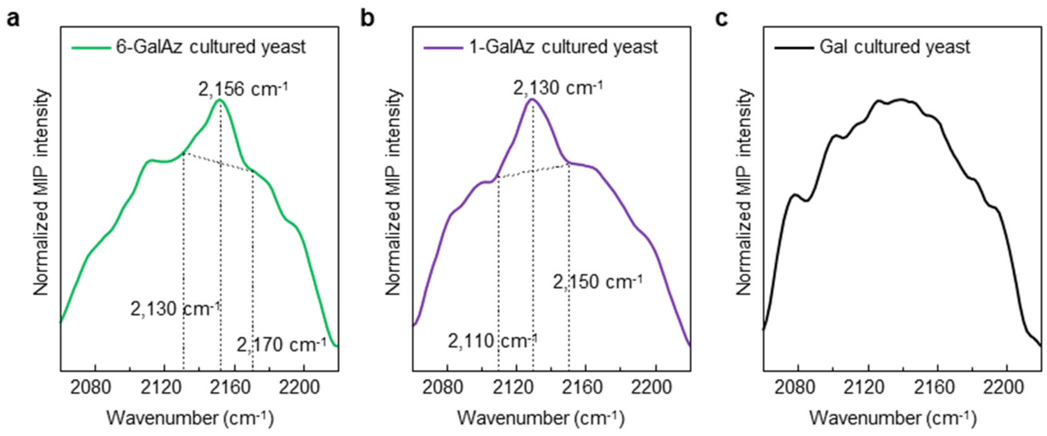


**Supplementary Fig. 14 Raw MIP spectra of yeast cells in the silent region.** MIP spectra acquired from (a) 6-GalAz, (b) 1-GalAz, and (c) Gal cultured yeast cells. The spectra are raw data without water background subtraction.

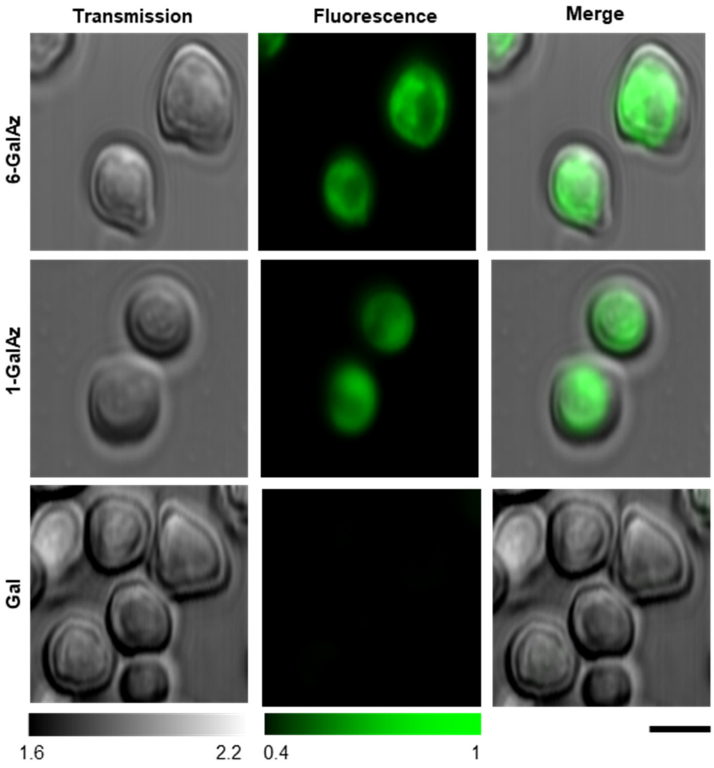


**Supplementary Fig. 15 Fluorescence images of single yeast cells cultured with 6-GalAz, 1-GalAz, Gal and underwent click reactions.** Scale bar: 5 µm.

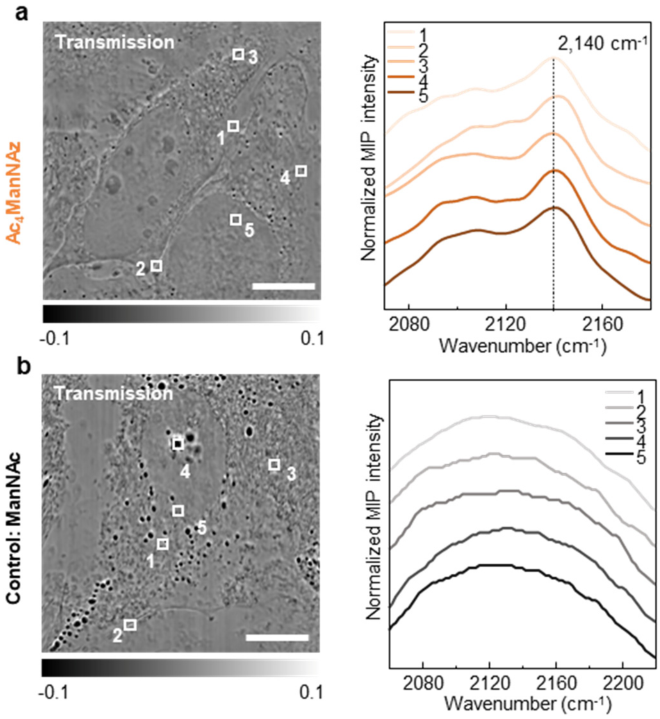


**Supplementary Fig. 16 Raw MIP spectra at indicated positions of HeLa cells in the silent region.** MIP spectra acquired from (a) Ac4ManNAz and (b) ManNAc treated HeLa cells shown in **Fig. 6d**. Scale bars: 20 µm. White boxes labeled in the transmission indicate the selected region (25 pixels) where the spectra were obtained. The spectra shown here are raw data without water background subtraction. Box 1, 2 in (a) and box 2 in (b) indicated the position of membranes.

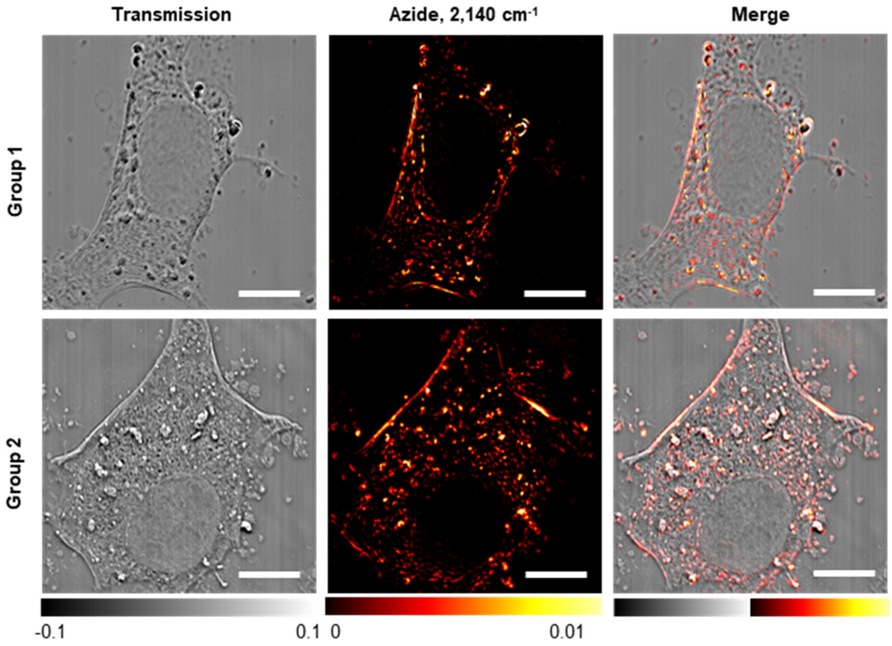


**Supplementary Fig. 17 MIP images of HeLa cells treated with $Ac_4ManNAz$.** The azide-conjugated carbohydrates on the cell surface and in the cytoplasm were clearly seen. Scale bars: 20 µm.

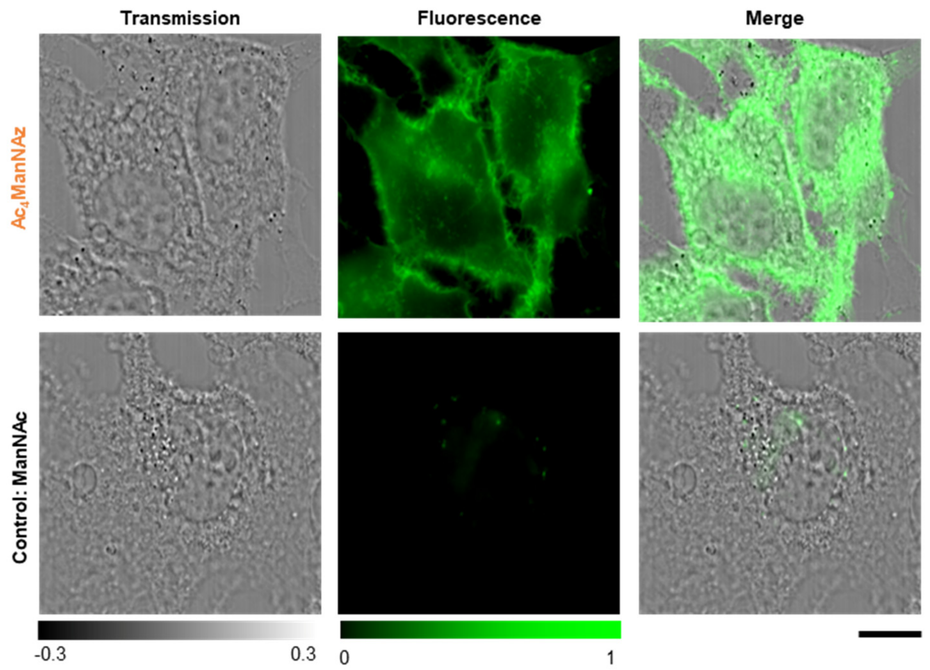


**Supplementary Fig. 18 Fluorescence images of HeLa cells treated with $Ac_4$ManNAz and ManNAc and underwent click reactions.** After 40-hour treatment of $Ac_4$ManNAz or ManNAc, the cells underwent click reactions with 1-hour incubation of 50 µM AF 488 DBCO. Scale bar: 20 µm.

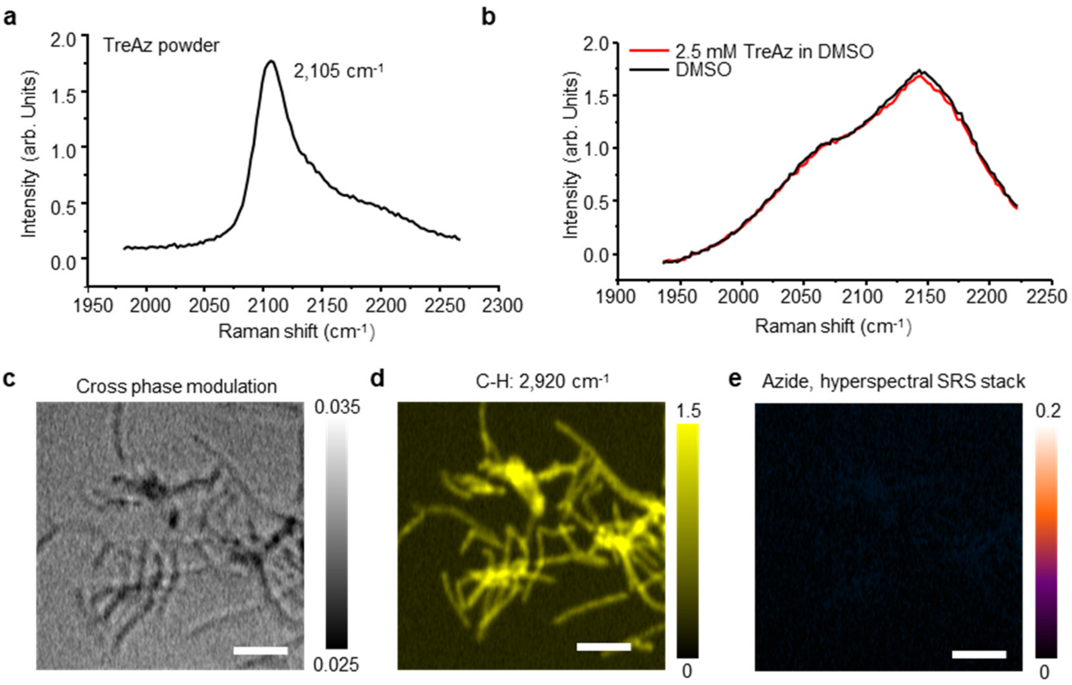


**Supplementary Fig. 19 Stimulated Raman scattering (SRS) spectroscopy and imaging of TreAz.** (a) SRS spectrum of pure TreAz powder, the azide peak is at 2,105 $cm^{-1}$. The pump beam wavelength was centered at 851 nm, and the power was 15 mW on sample. The probe beam wavelength was centered at 1,040 nm, and the power was 150 mW on sample. The pixel dwell time was 10 µs. (b) SRS spectra of 2.5 mM TreAz dissolved in DMSO and pure DMSO. The azide peak at the silent window is not detectable. The pump beam wavelength was centered at 851 nm, and the power was 15 mW on sample. The probe beam wavelength was centered at 1,040 nm, and the power was 150 mW on sample. The pixel dwell time was 100 µs. (c) Cross phase modulation imaging of *M. smeg* used as a reference. (d-e) Hyperspectral SRS imaging at the C-H channel (d) and the azide channel (e) of *M. smeg* treated with 50 µM TreAz for 26 hours. The C-H channel was recorded at 2,920 $cm^{-1}$. The Azide channel is a stack sum of 120-frame hyperspectral SRS image from 1,935.2 $cm^{-1}$ to 2,222.3 $cm^{-1}$. Scale bars: 5 µm. The pump beam wavelength was centered at 851 nm, and the power was 15 mW on sample. The probe beam wavelength was centered at 1,040 nm, and the power was 150 mW on sample. The pixel dwell time was 10 µs. Spectral focusing[10] was used to perform hyperspectral SRS imaging. A water objective with magnification of 60 and NA of 1.2 was used for focusing. An oil condenser with NA of 1.4 was used for signal collection.
